# Homeostatic control of synaptic rewiring in recurrent networks induces the formation of stable memory engrams

**DOI:** 10.1101/2020.03.27.011171

**Authors:** Júlia V. Gallinaro, Nebojša Gašparović, Stefan Rotter

## Abstract

Brain networks store new memories using functional and structural synaptic plasticity. Memory formation is generally attributed to Hebbian plasticity, while homeostatic plasticity is thought to have an ancillary role in stabilizing network dynamics. Here we report that homeostatic plasticity alone can also lead to the formation of stable memories. We analyze this phenomenon using a new theory of network remodeling, combined with numerical simulations of recurrent spiking neural networks that exhibit structural plasticity based on firing rate homeostasis. These networks are able to store repeatedly presented patterns and recall them upon the presentation of incomplete cues. Storage is fast, governed by the homeostatic drift. In contrast, forgetting is slow, driven by a diffusion process. Joint stimulation of neurons induces the growth of associative connections between them, leading to the formation of memory engrams. These memories are stored in a distributed fashion throughout connectivity matrix, and individual synaptic connections have only a small influence. Although memory-specific connections are increased in number, the total number of inputs and outputs of neurons undergo only small changes during stimulation. We find that homeostatic structural plasticity induces a specific type of “silent memories”, different from conventional attractor states.

## 1 Introduction

Memories are thought to be stored in the brain using cell assemblies that emerge through coordinated synaptic plasticity [53]. Cell assemblies with strong enough recurrent connections lead to bistable firing rates, which allows a network to encode memories as dynamic attractor states [39, 64]. If strong excitatory recurrent connections are counteracted by inhibitory plasticity, “silent” memories are formed [61, 47, 62]. In principle, assemblies can be generated by strengthening already existing synapses [39, 64], but potentially also by increasing connectivity among neurons. It has been shown that attractor networks can emerge through the creation of such neuronal clusters [38].

The creation of clusters through changes in connectivity between cells would require synaptic rewiring, or structural plasticity. Structural plasticity has been frequently reported in different areas of the brain, and sprouting and pruning of synaptic contacts was found to be often activity-dependent [29, 11, 40]. Sustained turnover of synapses, however, poses a severe challenge to the idea of memories being stored in synaptic connections [42]. Interestingly, recent theoretical work has shown that stable assemblies can be maintained despite ongoing synaptic rewiring [14, 16].

The formation of neuronal assemblies, or clusters, is traditionally attributed to Hebbian plasticity, driven by the correlation between pre- and postsynaptic neuronal activity on a certain time scale. For a typical Hebbian rule, a positive correlation in activity causes an increase in synaptic weight, which in turn increases the correlation between neuronal firing. This positive feedback cycle can result in unbounded growth, runaway activity and an essential dynamic instability of the network, if additional regulatory mechanisms are lacking. In fact, neuronal networks of the brain appear to employ homeostatic control mechanisms that regulate neuronal activity [59], and actively stabilize the firing rate of individual neurons at specific target levels [26, 55]. However, even though homeostatic mechanisms have been reported in experiments to operate on a range of different time scales, they seem to be too slow to trap the instabilities caused by Hebbian learning rules [65]. All things considered, it remains to be elucidated, what are the exact roles of Hebbian and homeostatic plasticity, and how these different processes interact to form cell assemblies in a robust and stable way [33].

Concerning the interplay between Hebbian and homeostatic plasticity, we have recently demonstrated by simulations that homeostatic structural plasticity alone can lead to the formation of assemblies of strongly interconnected neurons [18]. Moreover, we found that varying the strength of the stimulation and the fraction of stimulated neurons in combination with repetitive protocols can lead to even stronger assemblies [41]. In both papers, we used a structural plasticity model based on firing rate homeostasis, which had been used before to study synaptic rewiring linked with neurogenesis [8, 9], and the role of structural plasticity after focal stroke [5, 7] and after retinal lesion [4]. This model has also been used to study the emergence of criticality in developing networks [54] and other topological aspects of plastic networks [6]. Similarly to models with inhibitory plasticity [61, 47, 62], the memories formed in networks of this type represent a form of silent memory that is not in any obvious way reflected by neuronal activity.

The long-standing discussion about memory engrams in the brain has been revived recently. Researchers were able to identify and manipulate engrams [32], and to allocate memories to specific neurons during classical conditioning tasks [31]. These authors have also emphasized that an engram is not yet a memory, but merely the physical substrate of a potential memory in the brain [32]. Similar to the idea of a memory trace, it should provide the necessary conditions for a retrievable memory to emerge. Normally, the process of engram formation is thought to involve the strengthening of already existing synaptic connections. Here, we propose that new engrams could also be formed by a special form of synaptic clustering with increased synaptic connectivity among participating neurons, but the total number of incoming and outgoing synapses remaining approximately constant.

To demonstrate the feasibility of the idea, we performed numerical simulations of a classical conditioning task in a recurrent network with structural plasticity based on firing rate homeostasis. We were able to show that the cell assemblies formed share all characteristics of a memory engram. We further explored the properties of the formed engrams and developed a mean-field theory to explain the mechanisms of memory formation with homeostatic structural plasticity. We showed that these networks are able to effectively store repeatedly presented patterns. The formed engrams implement a special type of silent memory, which normally exists in a quiescent state and can be successfully retrieved using incomplete cues.

## 2 Results

### 2.1 Formation of memory engrams by homeostatic structural plasticity

We simulated a classical conditioning paradigm using a recurrent network. The network was composed of excitatory and inhibitory leaky integrate-and-fire neurons, and the excitatory-to-excitatory connections were subject to structural plasticity regulated by firing rate homeostasis. All neurons in the network received a constant background input in the form of independent Poisson spike trains. Three different non-overlapping subsets of neurons were sampled randomly from the network. The various stimuli considered here were conceived as increased external input to one of the specific ensembles, or combinations thereof. As the stimuli were arranged exactly as in behavioral experiments, we also adopted their terminology “unconditioned stimulus” (US) and “conditioned stimulus” (C1 and C2). The unconditioned response (UR) was conceived as the activity of a single readout neuron, which received input from the ensemble of excitatory neurons associated with the US (Figure 1A, top).

**Figure 1:**
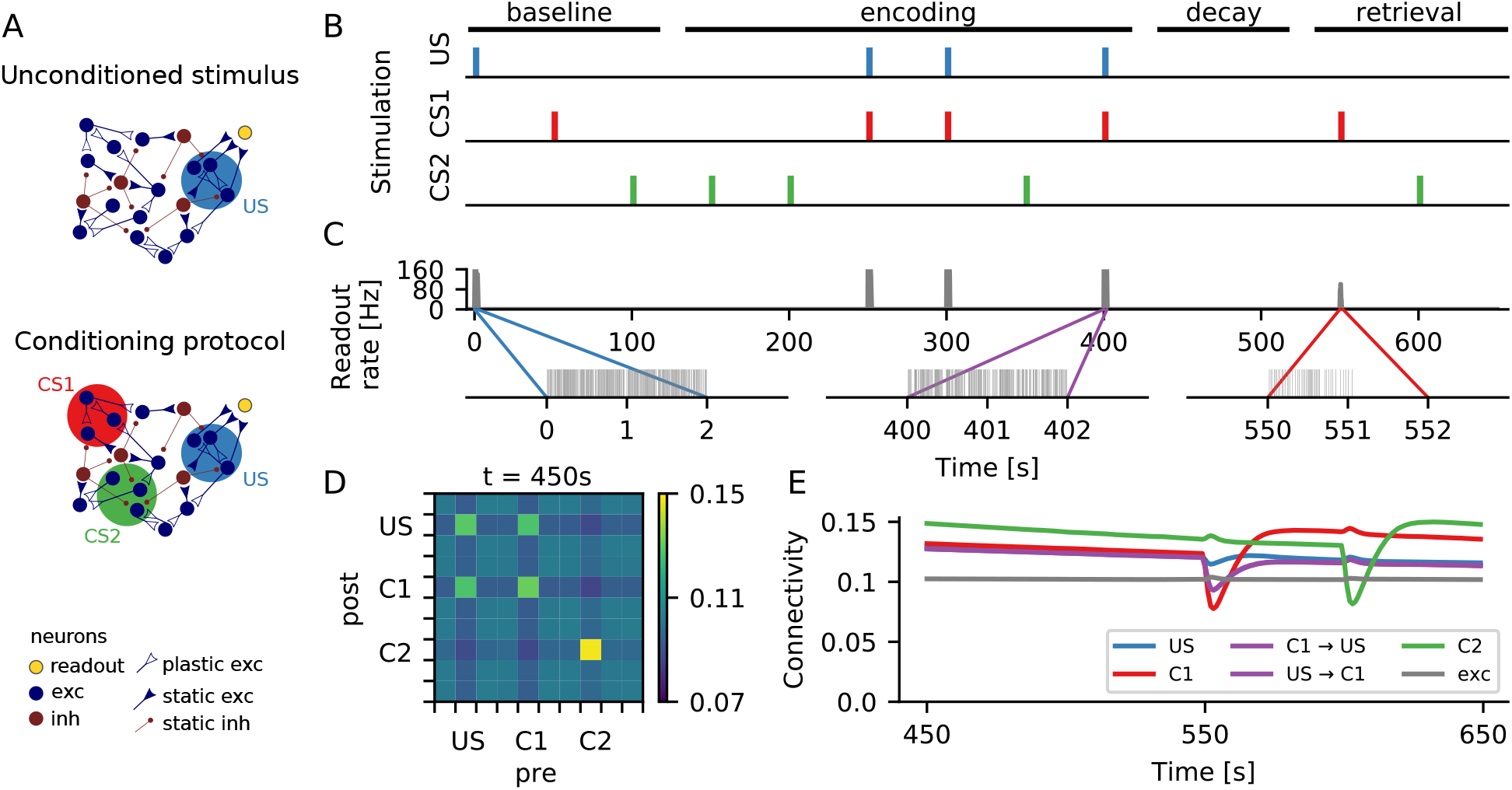
Formation of memory engrams in a neuronal network with homeostatic structural plasticity. (A) In a classical conditioning scenario, an unconditioned stimulus (US) was represented by a group of neurons that were connected to a readout neuron (yellow) via static synapses. The readout neuron spiked whenever neuronal activity representing the US was above a certain threshold (top). During a conditioning protocol, two other groups of neurons (CS1 and CS2) were chosen to represent two different conditioned stimuli (bottom). (B) During stimulation, the external input to a specific group of neurons was increased. The color marks indicate when each specific group was being stimulated. During the “encoding” phase, CS1 was always stimulated together with US, and CS2 was always stimulated alone. The time axis matches that of panel C (top). (C) Firing rate (top) and spike train (bottom) of the readout neuron. Before the paired stimulation (“baseline”), the readout neuron responded strongly only upon direct stimulation of the neuronal ensemble corresponding to the US. After the paired stimulation (“retrieval”), however, a presentation of C1 alone triggered a strong response of the readout neuron. This was not the case for a presentation of C2 alone. (D) Coarse grained connectivity matrix. Neurons are divided into 10 groups of 100 neurons each, shown is the average connectivity within each group. After encoding, the connectivity matrix indicates that engrams were formed, and we found enhanced connectivity within all three ensembles as a consequence of repeated stimulation. Bidirectional inter-connectivity across different engrams, however, is only observed for the pair C1-US that experienced paired stimulation. (E) Average connectivity as a function of time. The connectivity dynamics shows that engram identity was strengthened with each stimulus presentation, and that engrams decayed during unspecific external stimulation.

Figure 1B illustrates the protocol of the conditioning experiment simulated here. hlDuring a baseline period, the engrams representing US, C1 and C2 were stimulated once, one after the other. The stimulation consisted of increasing the rate of the background input by 40% for a period of 2 *mathrms.* In this phase, the activity of the US ensemble was high only upon direct stimulation (Figure 1B, middle). hlThe baseline period was followed by an encoding period, in which the C1 engram was always stimulated together with the US engram, while the C2 engram was always stimulated in isolation. Simultaneous stimulation of neurons in a recurrent network with homeostatic structural plasticity can lead to the formation of reinforced ensembles [18], which are strengthened by repetitive stimulation [41]. This is also what happened here: After the encoding period, each of the three neuronal ensembles had increased within-ensemble connectivity, as compared to baseline. Memory traces, or engrams, were formed (Figure 1D). Moreover, the US and C1 engrams also had higher bidirectional across-ensemble connectivity, representing an association between their corresponding memories.

Between encoding and retrieval, memory traces remained in a dormant state. Due to the homeostatic nature of network remodeling, the ongoing activity after encoding was very similar to the activity before encoding, but specific rewiring of input and output connections led to the formation of structural engrams. It turns out that these “silent memories” are quite persistent, as “forgetting” is much slower than “learning” them. In later sections, we will present a detailed analysis of this phenomenon. Any silent memory can be retrieved with a cue. In our case, this is a presentation of the conditioned stimulus. Stimulation of C1 alone, but not of C2 alone, triggered a conditioned response (Figure 1C) that was similar to the unconditioned response. Inevitably, stimulation of C1 and C2 during recall briefly destabilized the corresponding cell assemblies (seen as a drop in connectivity in Figure 1E), as homeostatic plasticity was still ongoing. The corresponding engrams then went through a reconsolidation period, during which the within-assembly connectivity grew even higher than before retrieval (Figure 1E, red and green). As a consequence, stored memories got stronger with each recall. Interestingly, as in our case the retrieval involved stimulation of either C1 or C2 alone, the connectivity between the US and C1 engrams decreased a bit after the recall (Figure 1E, purple).

Memories and associations were formed by changes in synaptic wiring, triggered by neuronal activity during the encoding period. They persisted in a dormant state and could be reactivated by a retrieval cue that reflected the activity experienced during encoding. This setting exactly characterizes a memory engram [32]. In the following sections, we will further characterize the process of formation of a single engram. We will also explore the nature and stability of the formed engram in more detail.

### 2.2 Engrams represent silent memories, not attractors

Learned engrams have a subtle influence on network activity. For a demonstration, we first grew a network under the influence of homeostatic structural plasticity (see 4.10.1). All neurons received a baseline stimulation in the form of independent Poisson spike trains. We then randomly selected an ensemble *E*_1_ of excitatory neurons and stimulated it repeatedly. Each stimulation cycle was comprised of a period of 150 s increased input to *E*_1_ and another 150 s relaxation period with no extra input. After 8 such stimulation cycles, the within-engram connectivity had increased to 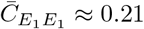. At this point, the ongoing activity of the network exhibited no apparent difference to the activity before engram encoding (Figure 2A). Due to the homeostatic nature of structural plasticity, neurons fired on average at their target rate, even though massive rewiring had led to higher within-engram connectivity. Looking closer, however, revealed a conspicuous change in the second-order properties of neuronal ensemble activity. We quantified this phenomenon using the overlap *m^μ^*. This quantity reflects the similarity of the neuron-by-neuron activity in a given time bin with a specified pattern, for example a pattern with all stimulated neurons being active and the remaining ones being silent (see Section 4.10.8 for a more detailed explanation of the concept). Figure 2B depicts the time-dependent (bin by bin) overlap of ongoing network activity with the engram *E*_1_ (*m*^*E*_1_^). It also shows the overlap with 10 different random ensembles *x* (*m^x^*), which are of the same size as *E*_1_ but have no neurons in common with it. The variance of *m*^*E*_1_^ is slightly larger than that of *m^x^* (Figure 2C). This indicates that the increased connectivity also increased the tendency of neurons belonging to the same learned engram to synchronize their activity, in comparison to other pairs of neurons. An increase in pairwise correlations within the engram, however, leads to increased fluctuations of the population activity [35], which also affects population measures such as the overlap used here.

**Figure 2.**
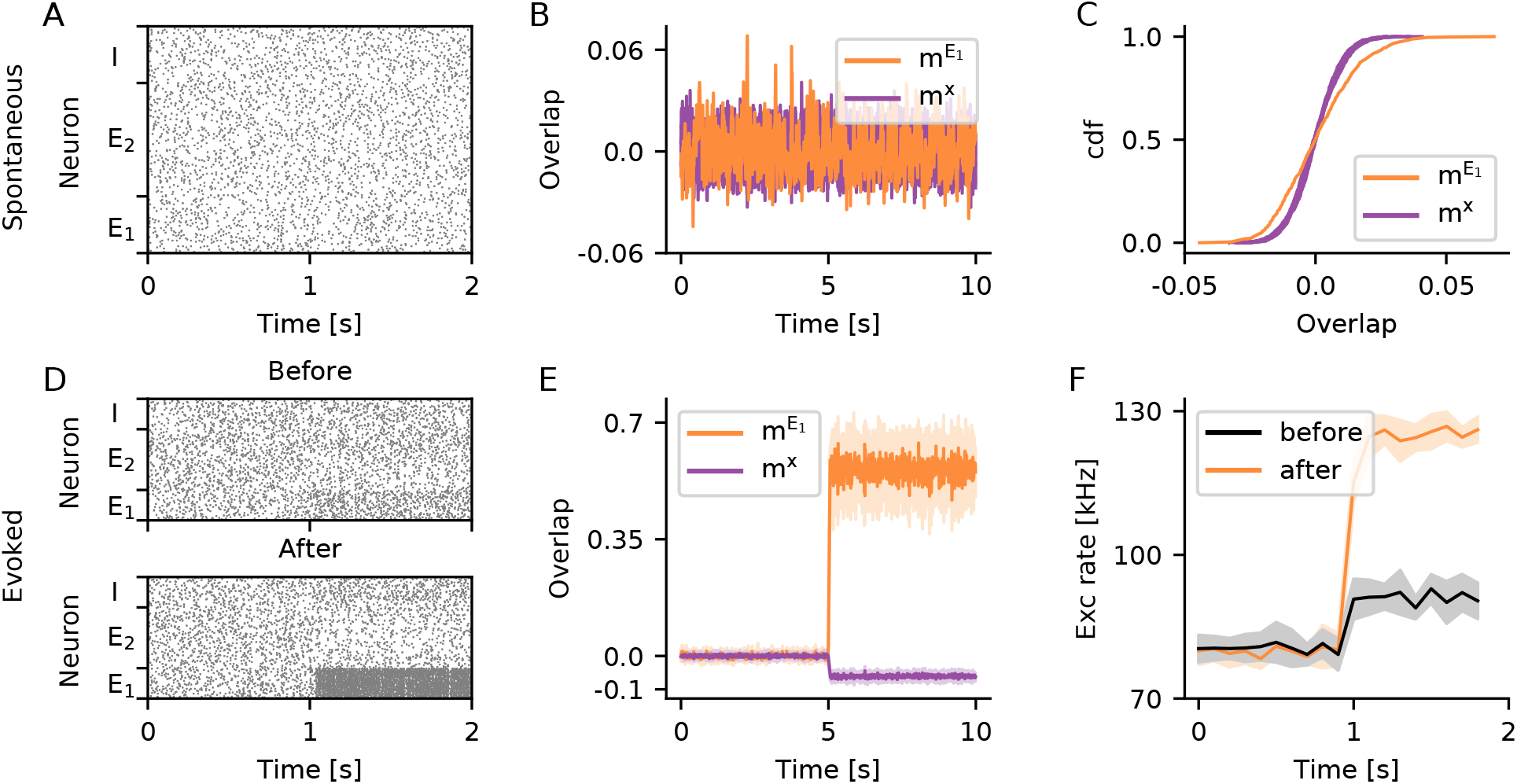
Silent memory based on structural engrams. (A–C) The ongoing activity of neurons belonging to the engram *E*_1_ can hardly be distinguished from the activity of the rest of the network. (A) Raster plot showing the spontaneous activity of 50 neurons randomly selected from *E*_1_, 100 neurons randomly selected from *E*_2_ but not belonging to *E*_1_, and 50 neurons randomly selected from the pool *I* of inhibitory neurons. (B) Overlap of ongoing network activity with the learned engram *E*_1_ (*m*^*E*_1_^, orange), and separately for 10 different random ensembles *x* disjoint with *E*_1_ (*m^x^*, purple). (C) Cumulative distribution of *m^μ^* shown in (B). (D–F) The activity evoked upon stimulation of *E*_1_ is higher, if the within-engram connectivity is large enough 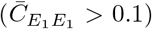 as a consequence of learning. (D) Same as A for evoked activity, the stimulation starts at *t* = 1 s. The neurons belonging to engram *E*_1_ are stimulated before (top, 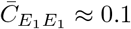) and after (bottom, 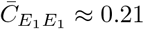) engram encoding. (E) Overlap with the learned engram (*m*^*E*_1_^, orange) and with random ensembles (*m^x^*, purple) during specific stimulation of engram *E*_1_. (F) Population rate of all excitatory neurons during stimulation of *E*_1_ before (black) and after (orange) engram encoding. (E, F) Solid line and shading depict mean and standard deviation across 10 independent simulation runs, respectively. In all panels, the bin size for calculating overlaps is 10 ms, and the bin size for calculating population rates is 100 ms.

During specific stimulation, the differences between the evoked activity of learned engrams and random ensembles were more pronounced. Figure 2D shows raster plots of network activity during stimulation of *E_1_* before and after the engram was encoded. The high recurrent connectivity within the *E*_1_ assembly after encoding amplified the effect of stimulation, leading to much higher firing rates of *E*_1_ neurons. This effect could even be seen in the population activity of all excitatory neurons in the network (Figure 2F). During stimulation, the increase in firing rate of engram neurons was accompanied by a suppression of activity of all other excitatory neurons not belonging to the engram. This was what underlied the conspicuous decrease in the overlap *m^x^* with random ensembles *x* during stimulation (Figure 2E).

How does the evoked response of an engram depend on the connectivity within? To answer this question, we looked into evoked activity at different points in time during stimulation. The within-engram connectivity increased with every stimulation cycle (Figure 3A), and so did the population activity of excitatory neurons during stimulation (Figure 3B). The in-degree of excitatory neurons, in contrast, was kept at a fixed level by the homeostatic controller, even after engram encoding (Figure 4C). This behavior was well captured by a simple mean-field firing rate model (grey line in Figure 3B), in which the within-engram connectivity was varied and all the remaining excitatory connections were adjusted to maintain a fixed in-degree of excitatory neurons.

**Figure 3.**
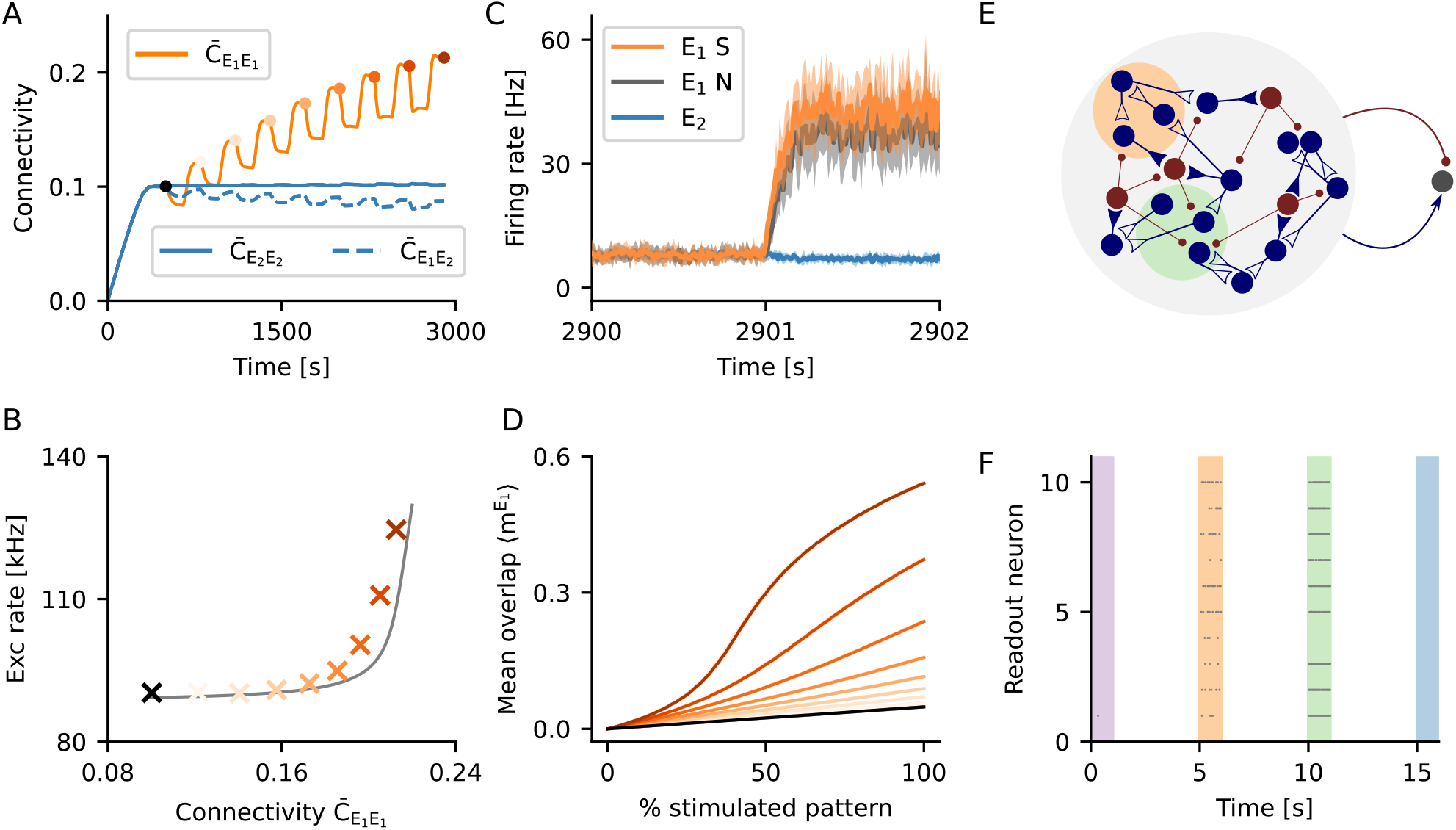
Evoked activity depended on the strength of a memory. (A) Starting from a random network grown under the influence of unstructured stimulation (black dot), we repeatedly stimulated the same ensemble of excitatory neurons *E*_1_ to eventually form an engram. Multiple stimulation cycles increased the rec urrent connectivity within the engram. (B) Population activity of all excitatory neurons upon stimulation of *E*_1_, for different levels of engram connectivity 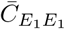. Crosses depict the population rate observed in a simulation. Colors indicate engram connectivity 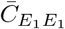, matching the colors used in panels (A) and (D). The grey line outlines the expectation from a simple mean-field theory. (C) Firing rate of *E*_1_ engram neurons upon stimulation of 50% its neurons. Shown is the mean firing rate of the stimulated engram neurons (*E*_1_*S*, orange), the mean firing rate of the non-stimulated engram neurons (*E*_1_*N*, grey), and the mean firing rate of excitatory neurons not belonging to the engram (*E*_2_, blue). Solid line and shading depict mean and standard deviation for 10 independent simulation runs, respectively. (D) Time-averaged overlap 〈*m*^*E*_1_^〉, for different fractions of *E*_1_ being stimulated. The recurrent nature of memory engrams enabled them to perform pattern completion. The degree of pattern completion depended monotonically on engram strength. (E) Two engrams (orange and green) were encoded in a network. Both engrams had a different strength with regard to their within-engram connectivity (green stronger than orange). A simple readout neuron received input from a random sample comprising 9 % of all excitatory and 9 % of all inhibitory neurons in the network. (F) Raster plot for the activity of 10 different readout neurons during the stimulation of learned engrams and random ensembles, respectively. Readout neurons were active when an encoded engram was stimulated (orange and green), and they generally responded with higher firing rates for stronger engrams (green). The activity of a readout neuron was low in absence of a stimulus (white), or upon stimulation of a random ensemble of neurons (purple and blue).

**Figure 4:**
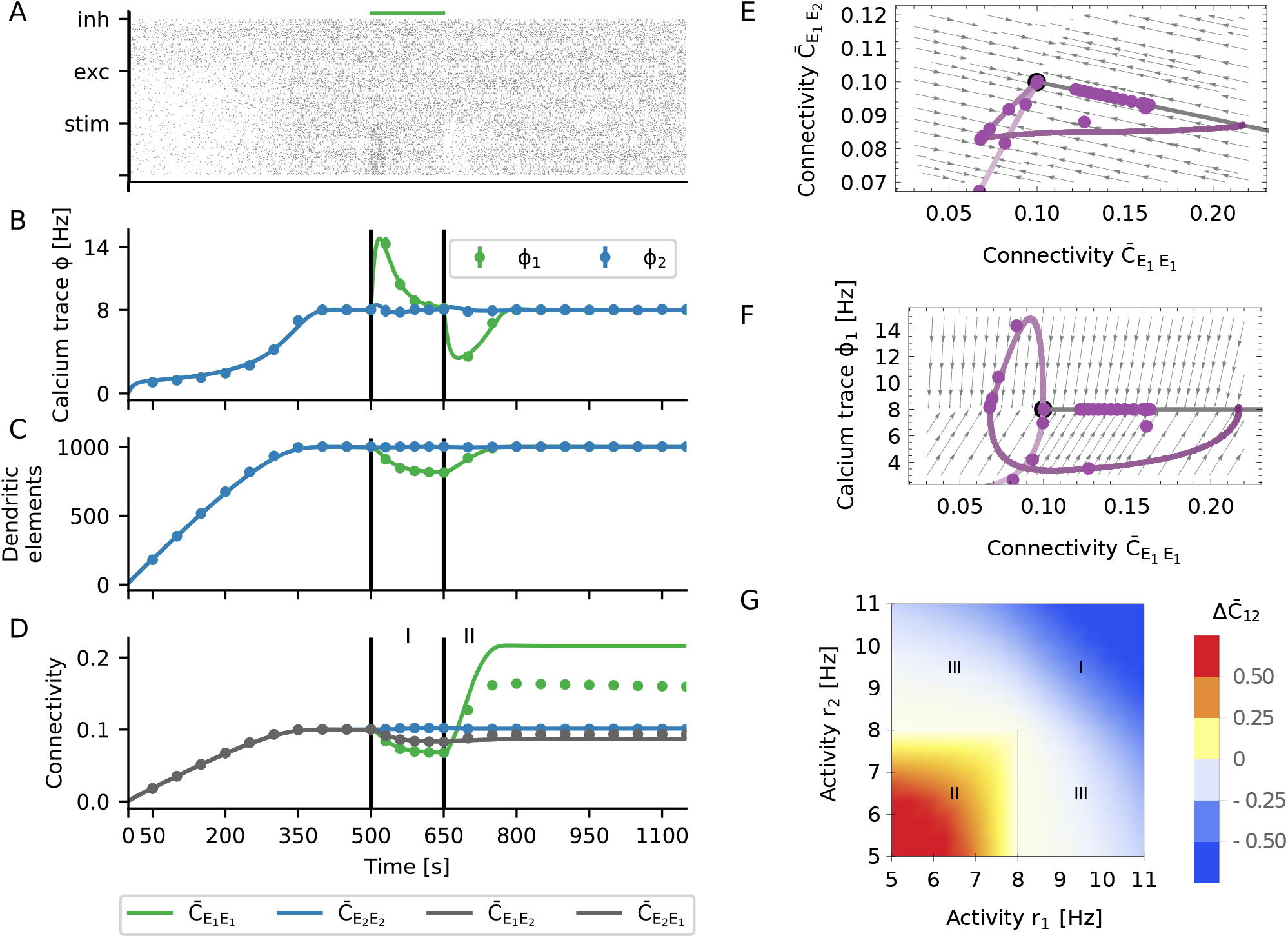
Hebbian properties emerge through the interaction of selective input and homeostatic control. (A) The activity of the neuronal network was subject to homeostatic control. For increased external input, it transiently responded with a higher firing rate. With a certain delay, the rate was down-regulated to the imposed set-point. When the stimulus was turned off, the network transiently responded with a lower firing rate, which was eventually up-regulated to the set-point again. The activity was generally characterized by irregular and asynchronous spike trains. (B) It was assumed that the intracellular calcium concentration followed the spiking dynamics, according to a first-order low-pass characteristic. Dots correspond to numerical simulations of the system, and solid lines reflect theoretical predictions from a mean-field model of dynamic network remodeling. (C) Dendritic elements (building blocks of synapses) were generated until an in-degree of *K*_ln_ = 1 000 was reached. This number slightly decreased during specific stimulation, but then recovered after the stimulus was removed. (D) Synaptic connectivity closely followed the dynamics of dendritic elements until the recovery phase, when the recurrent connectivity within the stimulated group *E*_1_ overshooted. (A–D) Black vertical lines indicate beginning and end of the stimulation. (E, F) Phase space representation of the activity. The purple lines are projections of the full, high-dimensional dynamics to different two-dimensional subspaces: (E) within-engram connectivity vs. across-ensemble connectivity and (F) within-engram connectivity vs. engram calcium trace. The dynamic flow was represented by the gray arrows. The steady state of the plastic network was characterized by a line attractor (thick gray line), defined by a fixed total in-degree and out-degree. The ensemble of stimulated neurons formed a stable engram, and the strength of the engram was encoded by its position on the line attractor. (G) The effective “instantaneous” learning rule for the expected connectivity between a pair of neurons is homeostatic in nature. It could also be viewed as an “inverse covariance rule” with baseline at the set-point of the homeostatic controller. The emerging Hebbian properties results from the more long-term combinatorial properties of rewiring across the whole network.

We were interested to know whether engrams performed pattern completion, and how this feature depended on the connectivity within. Following the formation of the engram *E*_1_ through 8 stimulation cycles (Figure 3A), we tested pattern completion by stimulating only 50% of the neurons in *E*_1_ (*E*_1_*S*), while the remaining neurons in *E*_1_ (*E*_1_*N*) were not directly stimulated. Figure 3C shows that, as expected, stimulation led to an increase in the activity of the directly stimulated neurons (*E*_1_*S*, orange). It also shows, however, a very similar increase of the activity of the remaining non-stimulated neurons (*E*_1_*N*, grey), indicating pattern completion. In order to quantitatively assess pattern completion at different levels of the connectivity within, we measured how the overlap of network activity within the engram, *m*^*E*_1_^, depended on partial stimulation. For an unstructured random network, *m*^*E*_1_^ increased at a certain rate with the fraction of stimulated neurons (Figure 3C, black line). We speak of “pattern completion”, if *m*^*E*_1_^ was systematically larger than in an unstructured random network. Figure 3C demonstrates clearly that the degree of pattern completion associated with a specific engram increased monotonically with the within-engram connectivity.

The evoked activity in learned engrams and in random ensembles of the same size is different. This feature can be taken as a marker for the existence of a stored memory. To demonstrate the potential of this idea, we employed a simple readout neuron for this task (Figure 3D). This neuron had the same properties as any other neuron in the network, and it received input from a random sample comprising a certain fraction (here 9 %) of all excitatory and the same fraction of all inhibitory neurons in the network. We encoded two engrams in the same network, one being slightly stronger (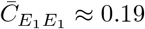, green) than the other one (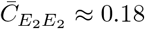, orange). We recorded the firing of the readout neuron during spontaneous activity, during specific stimulation of the engrams, and during stimulation of random ensembles of the same size. Figure 3E shows a raster plot of the activity of 10 different readout neurons, each of them sampling a different random subset of the network. With the parameters chosen here, the activity of a readout neuron was generally very low, except when a learned memory engram was stimulated. Due to the gradual increase in population activity with memory strength (Figure 3B), readout neurons responded with higher rates upon the stimulation of stronger engrams (green).

Homeostatic structural plasticity enables memories based on neuronal ensembles with increased within-ensemble connectivity, or engrams. Memories are acquired quickly and can persist for a long time. Moreover, the specific network configuration considered here admits a gradual response to the stimulation of an engram reflecting the strength of the memory. As we will show later, the engram connectivity lies on a line attractor, which turns into a slow manifold if fluctuations are taken into consideration. This configuration allows the network to simultaneously learn to recognize a stimulus (“Does the current stimulation correspond to a known memory?”) and to assess its confidence of the recognition (“How strong is the memory trace of this pattern?”). Such behavior would be absolutely impossible in a system that relies on bistable firing rates (attractors) to define engrams. Details of our analysis will be explained later in Section 2.4.

### 2.3 The mechanism of engram formation

We have shown how homeostatic structural plasticity created and maintained memory engrams, and we have then further elucidated the mechanisms underlying this process. We considered a minimal stimulation protocol [18] to study the encoding process for a single engram *E*_1_. We performed numerical simulations and developed a dynamical network theory to explain the emergence of associative (Hebbian) properties. In Figure 4, the results of numerical simulations are plotted together with the results of our theoretical analysis (see Section 4.6). Upon stimulation, the firing rate followed the typical homeostatic dynamics [60]. In the initial phase, the network stabilized at the target rate (Figure 4A). Upon external stimulation, it transiently responded with a higher firing rate. With a certain delay, the rate was down-regulated to the set-point. When the stimulus was turned off, the network transiently responded with a lower firing rate, which was eventually up-regulated to the set-point again.

Firing rate homeostasis relied on the intracellular calcium concentration *φ_i_*(*t*) of each neuron *i* (Figure 4D), which can be considered as a proxy for its firing rate. In our simulations, it was obtained as a low-pass filtered version of the spike train *S_i_*(*t*) of the neuron

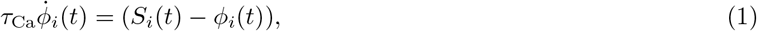

with time constant *τ_Ca_*. Each excitatory neuron *i* used its own calcium trace *φ_i_*(*t*) to control its number of synaptic elements. Deviations of the instantaneous firing rate (calcium concentration) *φ_i_*(*t*) from the target rate *v_i_* triggered either creation or deletion of elements according to

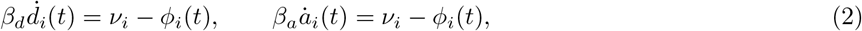

where *a_i_*(*t*) and *d_i_*(*t*) are the number of axonal and dendritic elements, respectively. The parameters *β_a_* and *β_d_* are the associated growth parameters (see Section 4.2 for more details). These equations are describing homeostatic control in our model. When activity is larger than the set-point, excitatory synapses are deleted. This tends to reduce activity. When activity is lower than the set-point, synaptic elements are created and new excitatory synapses are formed. This tends to increase activity. The number of elements are represented by continuous variables. A floor operation ⌊*x*⌋ is used to create the integers required for simulations.

During the initial growth phase, the number of elements increased to values corresponding to an in-degree *K*^in^ = *ϵN*, which was the number of excitatory inputs to the neuron that are necessary to sustain firing at the target rate (Figure 4D). Upon stimulation (during the learning phase), the number of connections was down-regulated due !05 to the transiently increased firing rate of neurons. After the stimulus was turned off, the activity returned to its 06 set-point. During the growing and learning phases, connectivity closely followed the dynamics of synaptic elements ‘07 (Figure 4D), and connectivity was proportional to the number of available synaptic elements (Figure 4D). After removal of the stimulus, however, in the reconsolidation phase, the recurrent connectivity within the stimulated group *E*_1_ was found to overshoot (Figure 4D). Then the average connectivity in the network returned to baseline (Figure 4C). While recurrent connectivity 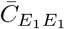 of the engram *E*_1_ increased, both the connectivity to the rest of network 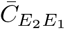 and from rest of the network 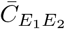 decreased, keeping the mean input to all neurons fixed. This indicates that although the network was globally subject to homeostatic control, local changes effectively exhibit associative features, as already pointed out in [18]. Our theoretical predictions generally match the simulations very well (Figure 4), with the exception that it predicted a larger overshoot. This discrepancy will be resolved in Section 2.4.

Deriving a theoretical framework of network remodeling (see Section 4.3) for the algorithm suggested by [4] posed a great challenge due to the large number of variables of both continuous (firing rates, calcium trace) and discrete (spike times, number of elements, connectivity, rewiring step) nature. The dimensionality of the system was effectively reduced by using a mean-field approach, which conveniently aggregated discrete counting variables into continuous averages (see Section 4.4 and Section 4.6 for more details of derivation).

Synaptic elements are accounted for by the number of free axonal *a*^+^(*t*) and free dendritic *d*^+^(*t*) elements, while the number of deleted elements is denoted by *a*^-^(*t*) and *d*^-^(*t*), respectively. Free axonal elements are paired with free dendritic elements in a completely random fashion to form synapses. The deletion of dendritic or axonal elements in neuron *i* automatically induces the deletion of incoming or outgoing synapses of that neuron, respectively. We employed a stochastic differential equation (Section 4.4) to describe the time evolution of connectivity 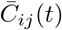 from neuron *j* to neuron *i*

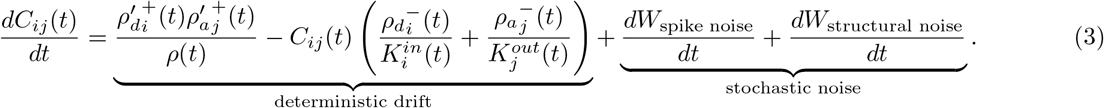

In this equation, 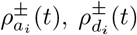 is the rate of creation/deletion of the axonal and dendritic elements of neuron i, respectively, and *ρ*(*t*) is the rate of creation of elements in the whole network. Note that *ρ*’^+^ is a corrected version of *ρ*^+^ (see Section 4.4 for details). The stochastic process described by Equation 3 decomposes into a deterministic drift process and a diffusive noise process. The noise process has two sources. The first source derives from the stochastic nature of the spike trains, and the second source is linked with the stochastic nature of axon-dendrite bonding. In this section, we ignore the noise and discuss only the deterministic part of the equation. This is equivalent to reducing the spiking dynamics to a firing rate model and, at the same time, treat connectivity in terms of its expectation values.

Stable steady-state solutions of the system described by Equation 3 represent a hyperplane in the space of connectivity (see Equation 17 in Section 4.7). These steady-state solutions are (random) network configurations with a fixed in-degree 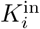 and out-degree 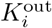 such that 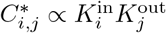. We will show that these states are attractive in Section 2.5 and Section 4.9. The hyperplane solution in our mean-field approach also generalizes to population variables (Section 4.6 and Section 4.7). In the case of only one engram stored in the network, there is the stimulated population and the “rest” of the network. In this case, the hyperplane from Equation 17 reduces to a line attractor of Equation 16 (Figure 4E and F, gray line). Memories are stored in the network in the following way: When a group of neurons is stimulated, the network diverges from the line attractor and takes a different path back during reconsolidation. The new position on the line encodes the strength of the memory. Stronger memories increase their connectivity within 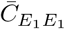 at expense of other connections. Furthermore, as the attractor is a skewed hyperplane in the space of connectivity, the memory is distributed across the whole neural network, and not only in recurrent connections among stimulated neurons in *E*_1_. As a reflection of this, other connectivity parameters (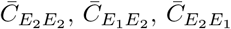, cf. Figure 4D) are also slightly changed.

To understand why changes in recurrent connectivity 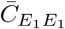 are associative, we note that the creation part of Equation 3 is actually a product 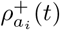 and 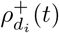, similar to a pre-post pair in a typical Hebbian rule. The main difference to a classical Hebbian rule is that only neurons firing below their target rate are creating new synapses. The effective rule is depicted in Figure 4G. Only neurons with free axonal or dendritic elements, respectively, can form new synapses, and those neurons are mostly the ones with low firing rates. The deletion part of Equation 3 depends linearly on 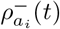 and 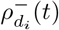, reducing to a simple multiplicative homeostasis. Upon excitatory stimulation, the homeostatic part is dominant and the number of synaptic elements decreases. Once the stimulus is terminated, the Hebbian part takes over, inducing a post-stimulation overshoot in connectivity. This leads to a peculiar bimodal dynamics of first decreasing connectivity and then overshooting, an important signature of this rule. We summarize this process in an effective rule

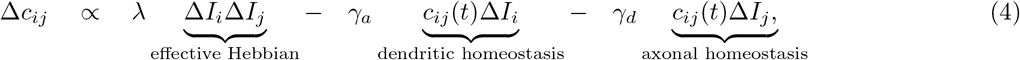

where Δ*I_i_* is the input perturbation of neuron *i*. The term Δ*I_i_*Δ*I_j_* is explicitly Hebbian with regard to input perturbations. Equation 4 only holds, however, if the stimulus is presented for a long enough time such that the calcium concentration tracks the change in activity and connectivity drops.

### 2.4 Fluctuation-driven decay of engrams

The qualitative aspects of memory formation have been explored in Section 2.3. Now we investigated the process of memory maintenance. A noticeable discrepancy between theory and numerical simulations was pointed out in Figure 4D. The overshoot is exaggerated and memories last forever. We found that this discrepancy was resolved when we take the spiking nature of neurons into account (Section 4.5).

Neurons use discrete spike trains 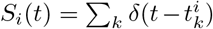 for signaling, and we conceived them here as stochastic point processes. We found that Gamma processes (Section 4.5) could reproduce the first two moments of the spike train statistics of the simulated networks with sufficient precision. The homeostatic controller in our model uses the trace of the calcium concentration *φ_i_*(*t*) as a proxy for the actual firing rate of the neuron. As the calcium trace *φ_i_*(*t*) is just a filtered version of the stochastic spike train *S_i_*(*t*), it is a stochastic process in its own respect. In Figure 5A we showed the stationary distribution of the time-dependent calcium concentration *φ_i_*(*t*). Apparently, a filtered Gamma process (purple line) provided a better fit to the simulated data than a filtered Poisson process (red line). The reason is that Gamma processes have an extra degree of freedom to match the irregularity of spike trains (coefficient of variation, here CV ≈ 0.7) as compared to Poisson processes (always CV = 1).

**Figure 5:**
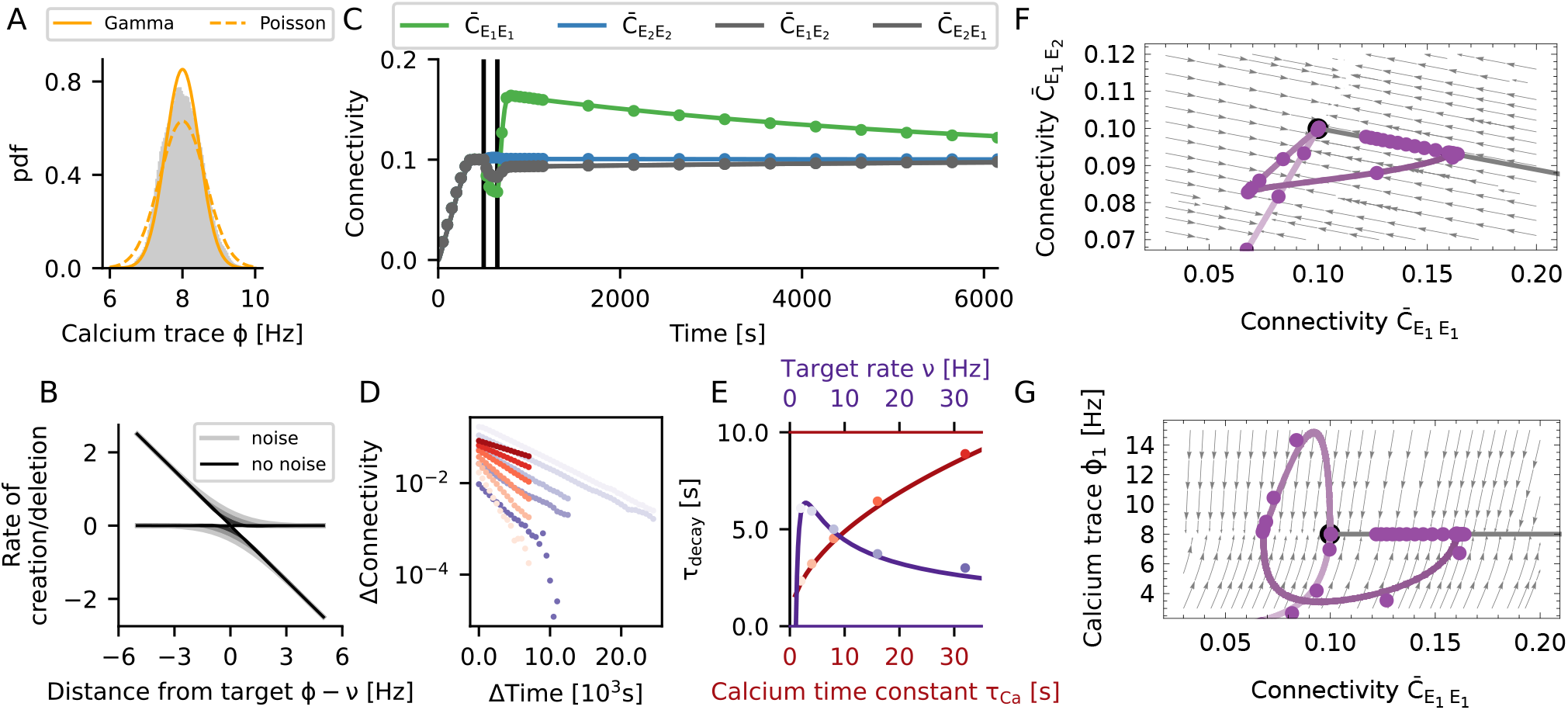
Noisy spiking induces fluctuations that lead to memory decay. (A) The gray histogram shows the distribution of calcium levels for a single neuron across 5 000 s of simulation. The yellow lines resulted from modeling the spike train as a Poisson process (dotted line) or a Gamma process (solid line), respectively. (B) The rate of creation or deletion of synaptic elements depends on the difference between the actual firing rate from the target rate (set-point), for different levels of spiking noise. The negative gain (slope) of the homeostatic controller in presence of noise is transformed into two separate processes of creation and deletion of synaptic elements. In the presence of noise (grey lines, lighter colors correspond to stronger noise), even when the firing rate is on target, residual fluctuations of the calcium signal induced a continuous rewiring of the network, corresponding to a diffusion process. (C) If noisy spiking and the associated diffusion was included in the model, our mean-field theory matched the simulation results very well. This concerns the initial decay, the overshoot and subsequent slow decay. Vertical lines indicate the beginning and the end of the stimulation. (D) Change in connectivity during the decay period, for different values of the calcium time constant (different shades of red, from light to dark *τ_Ca_* = 1, 2, 4, 8,16, 32 s, *ρ* = 8 Hz) and the target rate (different shades of purple, from light to dark *ρ* = 2,4, 8,16, 32 Hz, *τ_Ca_* = 10 s). We generally observed exponential relaxation as a consequence of a constant rewiring rate. (E) Time constant of the diffusive decay as a function of the calcium time constant and the target rate. Lines show our predictions from theory, and dots represent the values extracted from numerical simulations of plastic networks. The decay time *τ*_decay_ increases with 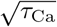. The memory was generally more stable for small target rates v, but collapsed for very small rates. This indicates an optimum for low firing rates, at about 3 Hz. (F, G) Same phase diagrams as shown in Figures 4E and F, but taking noise into consideration. (F) The spiking noise compromises the stability of the line attractor, which turns into a slow manifold. (G) The relaxation to the high-entropy connectivity configuration during the decay phase is indeed confined to a constant firing rate manifold.

The homeostatic controller (Equation 2) strived to stabilize *φ_i_*(*t*) at a fixed target value *v*, but *φ_i_*(*t*) fluctuates (Figure 5A) due to the random nature of the spiking. These fluctuations resulted in some degree of random creation and deletion of connections. Our theory (Section 4.5) reflected this aspect by an effective rule (Equation 12), which was obtained by averaging Equation 3 over the spiking noise (Figure 5B, red lines). The shape of this function indicates that connections were randomly created and deleted even when neurons were firing at their target rate. The larger the amplitude of the noise, the larger is the asymptotic variance of the process and the amount of spontaneous rewiring taking place.

We now extended our mean-field model of the rewiring process (see Section 2.3) to account for the spiking noise (see Section 4.5). According to the enhanced model, both the overshoot and the decay now matched very well with numerical simulations of plastic networks (Figure 5C). The decay of connectivity following its overshoot was exponential (Figure 2.4D), and Equation 3 revealed that the homeostasis was multiplicative and that the decay rate should be constant. The exponential nature of the decay is best understood by inspecting the phase space (Figure 5F and G). In terms of connectivity, the learning process was qualitatively the same as in the noise-less case (Figure 4E), where a small perturbation led to a fast relaxation to the line attractor (see Equation 16 in Section 4.7). In the presence of spiking noise (Figure 5F), however, the line attractor was deformed into a slow stochastic manifold (see Equation 18 in Section 4.8). The process of memory decay corresponded to a very slow movement toward the most entropic stable configuration compatible with firing rates clamped at their target value (Figure 5G). In our case, this led to a constraint on the in-degree *K*^ln^ = *ϵN_E_*, where *ϵ* is the mean equilibrium connectivity of excitatory neurons and *N_E_* is the number of excitatory neurons. In equilibrium, the connectivity matrix relaxed to a uniform connection probability 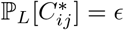. The realized connections, however, were constantly fluctuating. In every moment, the network had a configuration as for the Erdős-Rényi model. We refer to this state as “the” most entropic configuration. It will only change as a result of external stimulation. We summarize the memory decay process by the equation

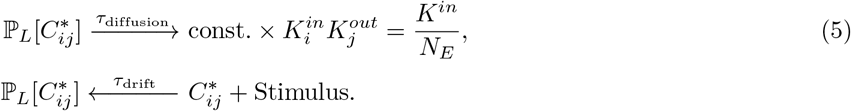

The fast “drift” process of relaxation back toward the slow manifold corresponds to the deterministic part of Equation 3. In contrast, the slow “diffusion” process of memory decay along the slow manifold corresponds to the stochastic part of Equation 3. In the equation above, *τ*_drlft_ represents the time scale of the fast drift process, and *τ*_dlffuslon_ is the time scale of the slow diffusion process.

The drift process is strongly non-linear, and its bandwidth is limited by the time constant of the calcium filter *τ*_Ca_, but also by the growth parameters of the dendritic elements *β_d_* and axonal elements *β_a_*. The diffusion process, on the other hand, is essentially constrained to the slow manifold of constant in-degree and firing rate (Section 4.8). Analytic calculations yield the relation

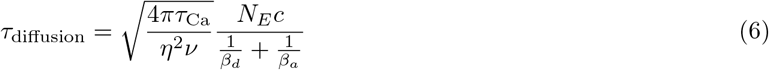

where *N_E_* is number of excitatory neurons, *c* is the average connectivity between excitatory neurons, and *η* is a correction factor to account for the reduced irregularity of spike trains as compared to a Poisson process. Both size and connectivity of the network increase the longevity of stored memories. Assuming that neurons rewire at a constant speed, it takes more time to rewire more elements. There is an interesting interference with the noise process, as memory longevity depends on the time constant of calcium in proportion to 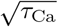 (Figure 5 E). On the other hand, the time scale of learning *τ*_drlft_ is limited by the low-pass characteristics of calcium, represented by *τ*_Ca_. Increasing *τ*_Ca_ leads to more persistent memories, but it also makes the learning slower. In the extreme case of *τ*_Ca_ → ∞ the system is unresponsive and exhibits no learning. As the calcium time constant, however, is finite, we don’t consider this possibility. We also find that the longevity of memories depends as 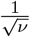 on the target rate. Therefore, making the target rate small enough should lead to very persistent memories. This path to very stable memories is not viable, though, because the average connectivity c implicitly changes with target rate. In Figure 5E longevity of the memory is depicted as a function of the target rate with corrected connectivity *c* = *c*(*ν*), and we find that for very small target rates memory longevity tends to zero instead. This suggests that there is an optimal range of target rates centered at a few spikes per second, fully consistent with experimental recordings from cortical neurons [10].

To summarize, the process of forming an engram (“learning”, see Section 2.3) exploits the properties of a line attractor. Taking spiking noise into account, the structure of the line attractor is deformed into a slow stochastic manifold. This still allows learning, but introduces controlled “forgetting” as a new feature. Forgetting is not necessarily an undesirable property of a memory system. In a dynamic environment, it might be an advantage for the organism to forget non-persistent or unimportant aspects of it. Homeostatic plasticity implements a mechanism of forgetting with an exponential time profile. However, psychological forgetting curves are often found to follow a power law [49]. The presented model cannot account for this directly, and additional mechanisms (e.g., replay, repeated stimulation, or alternative forms of plasticity) might be necessary to get this. Strong memories are sustained for a longer period, but they will eventually also be forgotten. In the framework of this model, the only way to keep memories forever is to repeat the corresponding stimulus from time to time, as illustrated in Figure 1D. If we think of the frequency of occurrence as a measure of the relevance of a stimulus, this implies that irrelevant memories decay but the relevant ones remain. Memories are stored in a way where the increase in connectivity within the engram is accompanied by a decrease of connectivity with non-engram neurons. This means that storage is linked with distributed changes in the connectivity matrix, instead of being stored in specific and localized synaptic connections [42]. Forgetting is reflected by a diffusion to the most entropic network configuration along the slow manifold. As a result, the system performs continuous inference from a persistent stream of information about the environment. Already stored memories are constantly refreshed in terms of a movement in directions away from the most entropic point of the slow manifold (novel memories define new directions), while diffusion pushes the system back to the most entropic configuration.

### 2.5 Network stability and constraints on growth parameters

So far, we have described the process of forming and maintaining memory engrams based on homeostatic structural plasticity. We have explained the mechanisms behind the striking associative properties of the system. Now, we will explore the limits of stability of networks with homeostatic structural plasticity and derive meaningful parameter regimes for a robust memory system. A homeostatic controller that operates on the basis of firing rates can be expected to be very stable by construction. Indeed we find that, whenever parameters are assigned meaningful values, the Jacobian 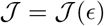 obtained by linearization of the system around the stable mean connectivity *ϵ* has only eigenvalues λ with non-positive real parts Re[λ] ≤ 0 (see Section 4.9). As a demonstration, we consider the real part of the two “most unstable” eigenvalues as a function of two relevant parameters, the dendritic/axonal growth parameter *β_d_* and the calcium time constant *τ*_Ca_ (Figure 6B, upper panel). The real part of these eigenvalues remains negative for any meaningful choice of time constants. It should be noted at this point, however, that the system under consideration is strongly non-linear, and linear stability alone does not guarantee global stability. We will discuss an interesting case of non-linear instability in Section 2.6. Oscillatory transients represent another potential issue in general control systems, and we will now explore the damped oscillatory phase of activity in more detail.

**Figure 6:**
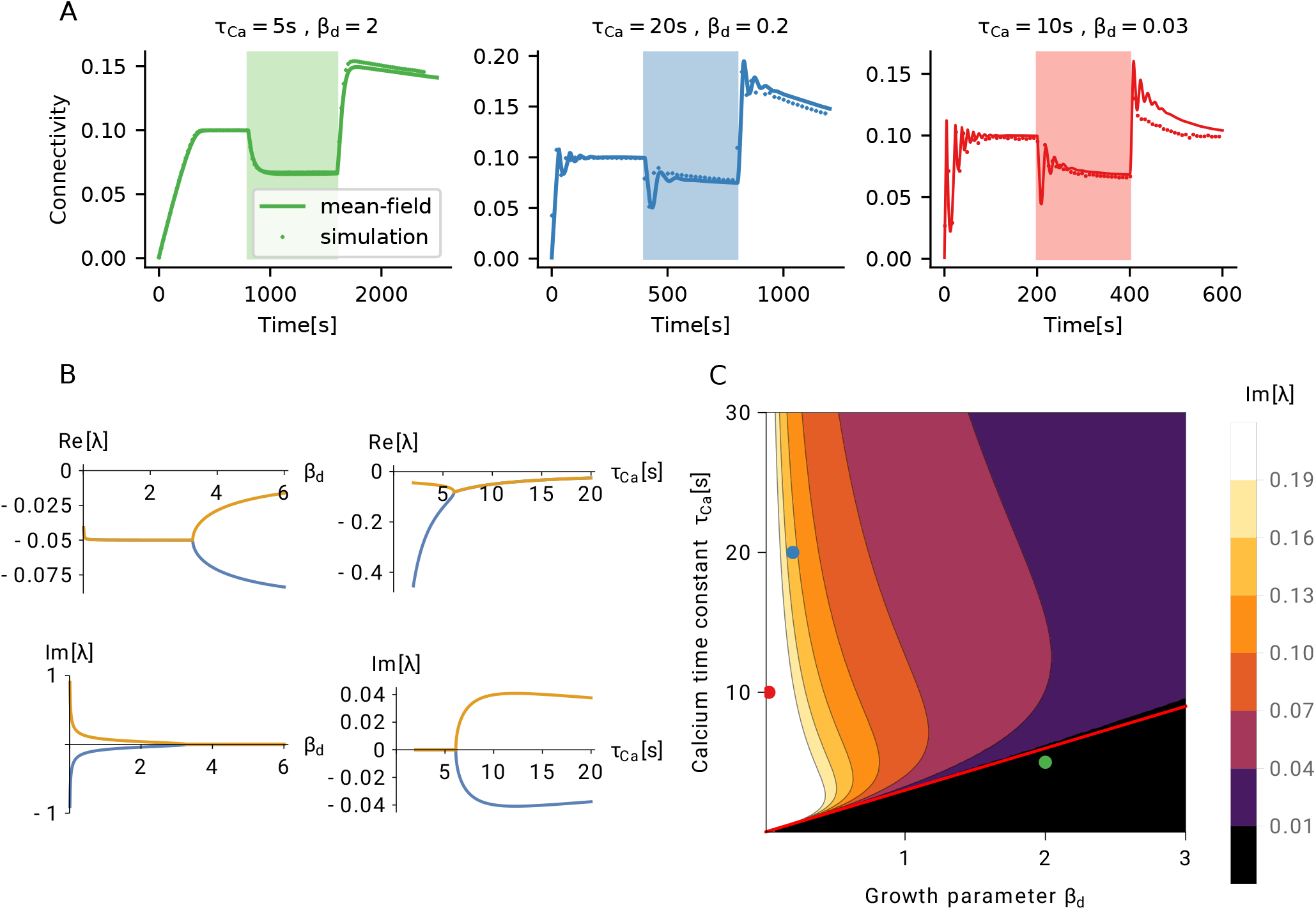
Linear stability of a network with homeostatic structural plasticity. (A) For a wide parameter regime, the structural evolution of the network has a single fixed point, which is also stable. Three typical types of homeostatic growth responses are depicted for this configuration: non-oscillatory (left), weakly oscillatory (middle), and strongly oscillatory (right) network remodeling. (B) All eigenvalues of the linearized system have a negative real part, for all values of the growth parameters of dendritic (axonal) elements *β_d_* and calcium *τ*_Ca_. In the case of fast synaptic elements (small *β_d_*) or slow calcium (large *τ*_Ca_), the system exhibits oscillatory responses. Shown are real parts and imaginary parts of the two “most unstable” eigenvalues, for different values of *β_d_* (left column, *τ*_Ca_ = 10 s) and *τ*_Ca_ (right column, *β_d_* = 2). (C) Phase diagram of the linear response. The black region below the red line indicates non-oscillatory responses, which corresponds to the configuration *τ*_Ca_ ≤ 3*β_d_* s. Dots indicate the parameter configurations shown in panel (A), with matching colors.

In Figure 6A we depict three typical cases of homeostatic growth responses: non-oscillatory (left, green), weakly oscillatory (middle, blue), and strongly oscillatory (right, red) network remodeling. The imaginary parts of the two eigenvalues shown in Figure 6B (bottom left), which are actually responsible for the oscillations, become non-zero when the dendritic and axonal growth parameters are too small (for other parameters, see Section 2.3), and both creation and deletion of elements are too fast. Oscillations occur, on the other hand, for large values of the calcium time constant *τ*_Ca_ (Figure 6B, bottom right). The system oscillates, if the low-pass filter is too slow as compared to the turnover of synaptic elements. The combination of *β_d_* and *τ*_Ca_ that leads to the onset of oscillations can be derived from the condition Im[λ] = 0. We can further exploit the fact that two oscillatory eigenvalues are complex conjugates of each other, and that the imaginary part is zero, when the real part bifurcates.

To elucidate the relative importance of the two parameters *β_d_* and *τ*_Ca_, we now explore how they together contribute to the emergence of oscillations (Figure 6C). The effect of parameters on oscillations is a combination of the two mechanisms discussed above: low-pass filtering and agility of control. We use the bifurcation of the real part of the least stable eigenvalues as a criterion for the emergence of Im[λ] = 0, which yields the boundary between the oscillatory and the non-oscillatory region (Figure 6C, red line). The black region of the phase diagram corresponds to a simple fixed point with no oscillations, the green point corresponds to the case shown in Figure 6A, left. From Figure 6B, bottom, we conclude that the fastest oscillations are created when *β_d_* is very small, as the oscillation frequency then exhibits a 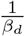 asymptotic dependence. The calcium dependence is a slowly changing function. The specific case indicated by a red point in Figure 6C corresponds to the dynamics shown in Figure 6A, right. Although intermediate parameter values result in damped oscillations (Figure 6C, blue dot; Figure 6A, middle), its amplitude remains relatively small. This link between two parameters can be used to predict a meaningful range of values for *β_d_*. Experimentally reported values for *τ*_Ca_ are combined with the heuristic of not exhibiting strong oscillations.

The analysis outlined in the previous paragraph clearly suggests that, in order to avoid excessive oscillations, the calcium signal (a proxy for neuronal activity) has to be faster than the process which creates elements. Oscillations in network growth are completely suppressed, if it is at least tree times faster. Strong oscillations can compromise non-linear stability, as we will show in Section 2.6. A combination of parameters that leads to oscillations could also imply a loss of stability for certain stimuli, for reasons explained later. We assume that the calcium time constant is in the range between 1 s and 10 s, in line with the values reported for somatic calcium transients in experiments [23, 24, 27, 67]. This indicates that homeostatic structural plasticity should not use element growth parameters smaller than around 0.4. Faster learning must be based on other types of synaptic plasticity (e.g. spike-timing dependent plasticity, or fast synaptic scaling).

We use this analysis framework now to compute turnover rates (TOR) and compare them to the values typically found in experiments. In Sections 2.3 and 2.4 we use a calcium time constant of 10 s and a growth parameters for synaptic elements of *β_a_* = *β_d_* = 2, which results in TOR of around 18% per day (see Methods). Interestingly, [56] measured TOR in the barrel cortex of young mice and found TOR values of around 20 % per day. After sensory deprivation, the TOR increased to a maximum of around 30 % per day in the barrel cortex (but not elsewhere). In our model, stimulus-dependent rewiring is strongest in the directly stimulated engram *E*_1_ (Figure 4C). This particular ensemble rewires close to 25 % of its dendritic elements per stimulation cycle. This very large turnover is a result of our experimental design involving a very strong stimulus. In principle, we could choose weaker stimuli, but then we would need many more encoding episodes, as described in [41]. The main results of the paper, however, would remain the same.

### 2.6 Loss of control leads to bursts of high activity

A network the connectivity of which is subject to homeostatic regulation generally exhibits robust linear stability around the fixed point of connectivity *ϵ*, as explained in detail in Section 2.5 and Section 4.9. But what happens, if the system is forced far away from its equilibrium? To illustrate the new phenomena arising, we repeat the stimulation protocol described in Section 2.3 with one stimulated ensemble *E*_1_. However, we now increase both the strength and the duration of the stimulation (Figure 7A). The network behaves as before during the growth and the stimulation phase (Figure 7A, upper panel), but during the reconsolidation phase connectivity gets out of control. New recurrent synapses are formed at a very high rate until excessive feedback of activity triggered by input from the non-stimulated ensemble causes an explosion of firing rates (Figure 7A bottom panel). The homeostatic response of the network to such seizure-like activity can only be a brisk decrease in recurrent connectivity. As a consequence, the activity *r*_E_1__ quickly drops to zero and the deregulated growth cycle starts all over.

**Figure 7:**
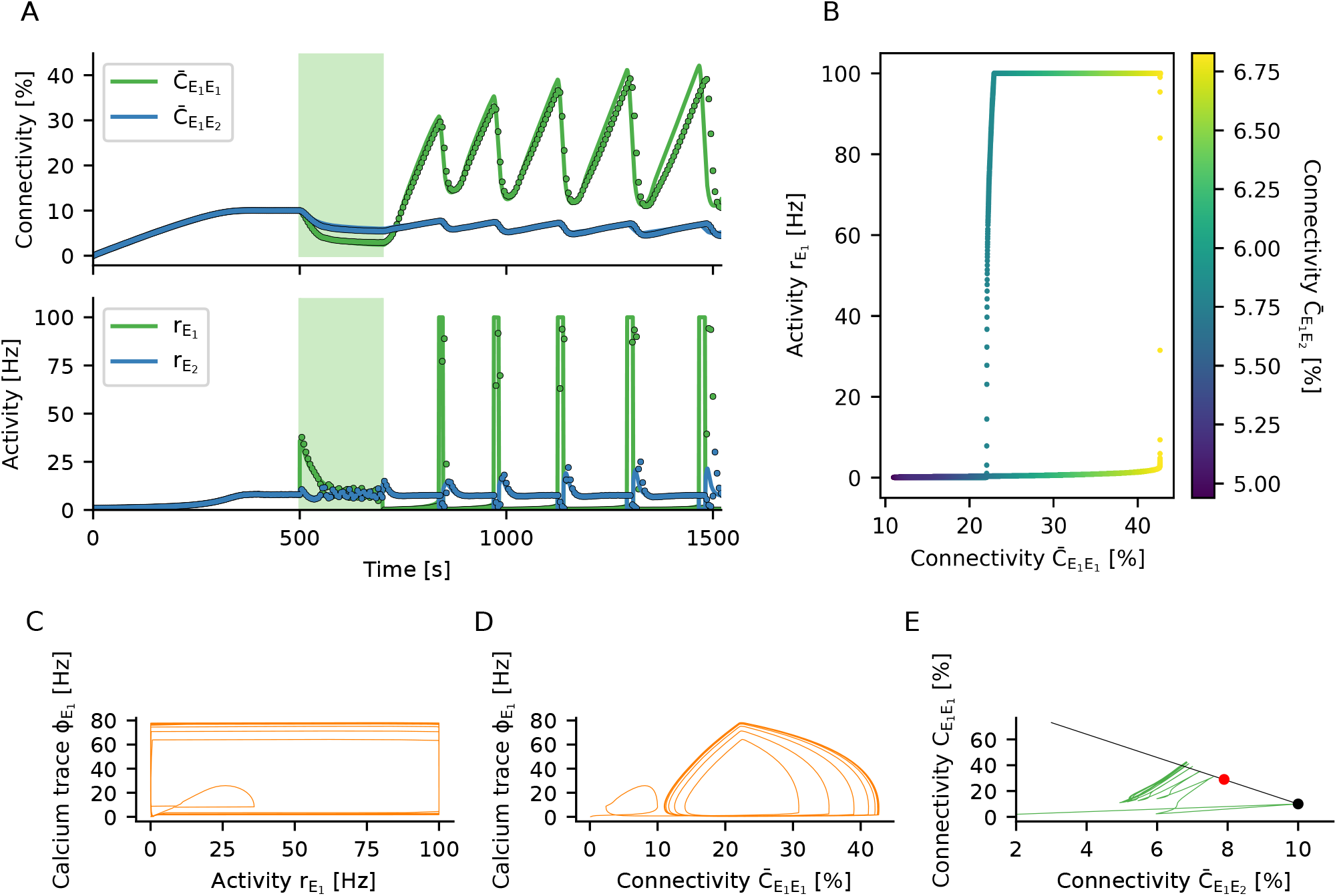
Non-linear stability of a network with homeostatic structural plasticity. (A) The engram *E*_1_ is stimulated with a very strong external input. As the homeostatic response triggers excessive pruning of recurrent connections, the population *E*_1_ is completely silenced after the stimulus is turned off. This, in turn, initiates a strong compensatory overshoot of connectivity and consecutive runaway population activity. The dots with corresponding color show the results of a plastic network simulation, and the solid lines indicate the corresponding predictions from our theory. The theoretical instantaneous firing rate is clipped at 100 Hz. (B) The network settles in a limit cycle of connectivity dynamics. The hysteresis-like behavior is caused by the faster growth of within-engram connectivity 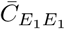 as compared to connectivity from the non-engram ensemble 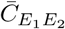. During the initial phase of the cycle, the increase of 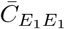 has no effect on the activity of population *E*_1_ yet, as its neurons are not active. Only when the input from population *E*_2_ through 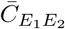 gets large enough, the rate *r*_*E*_1__ becomes non-zero and rises to very high values quickly due to already large recurrent 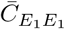 connectivity. (C) The calcium signal φ adds an additional delay to the cycle. (D) This leads to smoother trajectories when scattering calcium concentration against connectivity. (E) Connectivity within the stimulated group 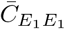 plotted against input connectivity from the non-engram population 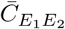. The black line shows configurations with constant in-degree, of which the black dot represents the most entropic one. The red dot corresponds to critical connectivity, beyond which the limit cycle behavior is triggered. The limit cycle transients in connectivity space are orthogonal to the line attractor, indicating that the total in-degree is oscillating and no homeostatic equilibrium can be established.

We explore the mechanism underlying this runaway process by plotting the long-term dynamics in a phase plane spanned by recurrent connectivity 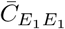 and the activity of the engram *r*_*E*_1__ (Figure 7B). A special type of limit cycle emerges, and we can track it using the input connectivity from the rest of the network to the learned engram (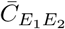, see Figure 7B). The cycle is started when the engram *E*_1_ is stimulated with a very strong external input. As the homeostatic response of the network triggers excessive pruning of its recurrent connections, the population *E*_1_ is completely silenced, *r*_*E*_1__ = 0, after the stimulus has been turned off. Then, homeostatic plasticity sets in and tries to compensate the activity below target by increasing the recurrent excitatory input to the engram *E*_1_. The growth of intra-ensemble connectivity 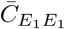 is faster than the changes in inter-ensemble connectivity 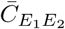, as the growth rate of 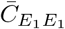 is quadratic in the rate *r*_*E*_1__ (Figure 4G), but 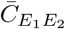 depends only linearly on it. However, while there are no recurrent spikes, *r*_*E*_1__ = 0, the increase in intra-ensemble connectivity cannot restore the activity to its target value. As soon as input from the rest of the network via 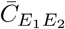 is strong enough to increase the rate *r*_*E*_1__ to non-zero values, engram neurons very quickly increase their own rate by activating recurrent connectivity 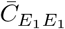. At this point, however, the network has entered a state in which the population activity is bistable, and there is a transition to high activity state, coinciding with a pronounced outbreak of population activity *r*_*E*_1__. The increase in rate is then immediately counteracted by the homeostatic controller. Due to the seizure-like activity burst, a high amount of calcium is accumulated in all participating cells. As a consequence, neurons delete many excitatory connections, and the firing rates are driven back to zero. This hysteresis-like cycle of events is repeated over and over again (Figure 7B), even if the stimulus has meanwhile been turned off. The period of the limit cycle is strongly influenced by the calcium variable, which lags behind activity (Figure 7C). Replacing activity *r*_*E*_1__ by recurrent connectivity 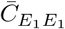, a somewhat smoother picture emerges (Figure 7D).

Two aspects are important for the emergence of the limit cycle. Firstly, the specific relation between the time scales of calcium and synaptic elements gives rise to different types of instabilities (see Figure 6 and Figure 7C and D). Secondly, the rates of creation and deletion of elements do not have the same bounds. While the rate with which elements are created *ρ*^+^ is limited by 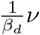, the rate of deletion *ρ*^-^ is limited by 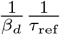. This peculiar asymmetry causes the observed brisk decrease in connectivity after an extreme seizure-like burst of activity. An appropriate choice of the calcium time constant, in combination with a strict limit on the rate of deletion, might lead to a system without the (pathological) limit cycle behavior observed in simulations. Finally, we have derived a criterion for bursts of population activity to arise, related to the loss of stability due to excessive recurrent connectivity (see Section 4.9, Equation 22). Indeed, a network with fixed in-degrees becomes dynamically unstable, if the connectivity of subpopulation *E*_1_ exceeds the critical value 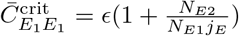, which for our parameters is at about 29%. The neurons comprising the engram *E*_1_ receive too much recurrent input (Figure 7E), and the balance of excitation and inhibition brakes. In this configuration, only one attractive fixed point exists for high firing rates, and a population burst is inevitable. In simulations, the stochastic nature of the system tends to elicit population bursts even earlier, at about 22% connectivity in our hands. We conclude that 22–29% connectivity is a region of bi-stability, with two attractive fixed points coexisting. Early during limit cycle development, the total in-degree is less than *ϵN* (the connectivity is in the region below the black line in Figure 7E), and the excitation-inhibition balance is broken by positive feedback. Later, the limit cycle settles into a configuration, where the total in-degree exceeds *ϵN* (the black line is crossed from below in Figure 4E). We have shown before that stable learning leads to silent memories in the network (Section 2.1 and 2.2), but in the case discussed here, sustained high activity is at odds with stable homeostatic control of network growth.

## 3 Discussion

We have demonstrated by numerical simulations and by mathematical analysis that structural plasticity controlled by firing rate homeostasis can implement a memory system based on the emergence and the decay of engrams. Input patterns are defined by a stimulation of the corresponding ensemble of neurons in a recurrent network. Presenting two patterns concurrently leads to their association by newly formed synaptic connections between the involved neurons. This mechanism can be used, among other things, to effectively implement classical conditioning. The memories are stored in network connectivity in a distributed fashion, defined by the engram as a whole and not by isolated individual connections. The memories are dynamic. They decay if previously learned stimuli are no longer presented, but they get stronger with every single recall. The memory is not affecting the firing rates during spontaneous activity, but even weak memory traces can be identified by the correlation of activity. Memories become visible as a firing rate increase of a specific pattern upon external stimulation, though. The embedding network is able to perform pattern completion, if a partial cue is presented. Finally, we have devised a simple recognition memory mechanism, in which downstream neurons respond with a higher firing rate, if any of the previously learned patterns is stimulated.

Memory engrams emerge because the homeostatic rule acts as an effective Hebbian rule with associative properties. This unexpected behavior is achieved by an interaction between the temporal dynamics of homeostatic control and a network-wide distributed formation of synapses. Memory formation is a fast process, exploiting degrees of freedom orthogonal to a line attractor while it reacts to the stimulus, and storing memories as positions on the line attractor. The spiking of neurons introduces fluctuations, which lead to the decay of memory on a slow time scale through diffusion along the line attractor. In absence of specific stimulation, the network slowly relaxes to the most entropic configuration of uniform connectivity across all pairs of neurons. In contrast, multiple repetitions of a stimulus push the system to states of lower entropy, corresponding to stronger memories. The dynamics of homeostatic networks is, by construction, very robust for a wide parameter range. Instabilities occur when the time scales of creating and deleting synaptic elements are much smaller than the time scales of the calcium trace, which feeds the homeostatic controller. Under these conditions, the network displays oscillations of fast decaying amplitudes, but it remains linearly stable. Stability is lost, though, when the stimulus is too strong. In this case, the compensatory forces lead to a limit cycle dynamics with pathologically large amplitudes.

Experiments involving engram manipulation have increased our current knowledge about this type of memory [32], and some recent findings are actually in accordance with our model. For example, memory re-consolidation was disrupted if a protein synthesis inhibitor was administered immediately after the retrieval cue during an auditory fear conditioning experiment [45]. In our model, engram connectivity initially decreases upon stimulation, and memories are shortly destabilized and consolidated again after every retrieval. Interfering with plasticity during or after retrieval could, therefore, also lead to active forgetting. Our model also predicts a decreased connectivity following cue retrieval, which should be visible if the disruption is imposed at the right point in time. We are, however, not aware of any experiments demonstrating this decreased connectivity. It is conceivable, therefore, that additional mechanisms exist in biological networks that actively counteract such decay. Another example of experiments that are in accordance to our model refers to excitability and engram allocation. Recent experiments have shown that neurons are more likely to be allocated to an engram, if they are more excitable before stimulation [63, 66]. In our model, more excitable neurons would fire more during stimulation, making them more likely to become part of an engram as a result of the increased synaptic turnover. Moreover, our analysis of the model suggests that decreasing excitability of some neurons soon after stimulation should also increase the likelihood that they become part of the engram. Further research with our model of homeostatic engram formation might include even more specific predictions for comparison with experiments involving engram manipulation. This would also help to better characterize and understand the process of engram formation in the brain.

The exact rules governing the sprouting and pruning of spines and boutons, and how these depend on neuronal activity, are unknown. While some studies show a constant turnover of spines, but fixed spine density [30], [56], other studies report either an increased spine density after stimulation in LTP protocols [43], [57], [1], or a loss of spines after persistent depolarization [44], [13], see [15] for a review on activity dependent structural plasticity. These different results might seem contradictory at first, but our model suggests that they could all be related. This is because the specific connectivity within the stimulated population undergoes a bimodal change over time (Figure 4 D), and whether one encounters an increase or a decrease in spine density would depend on when the measurements were performed. More specifically, our model predicts that the overall spine density remains unchanged before and after stimulation (Figure 4C), notwithstanding a small temporary decrease during stimulation. Most importantly, however, our model emphasizes a reallocation of synapses, resulting in higher connectivity between stimulated neurons and reduced connectivity to non-stimulated neurons. This means that the overall number of spines is the same before and after stimulation (same number of dendritic elements before and after stimulation on Figure 4). However, the number of spines connecting to other stimulated neurons is higher after stimulation (green line on Figure 4D) and the number of spines connecting to non-stimulated neurons is smaller (grey lines on Figure 4D). Demonstrating such specific effects experimentally would require to establish for each synapse separately, whether the presynaptic and the postsynaptic neuron was stimulated, or not. To our knowledge, there is currently no experimental study available reporting such labeled connectivity data. As the model is inherently stochastic, it is not necessary that a synapse is deleted each time their presynaptic or postsynaptic neurons are stimulated. Instead, random deletions between any pair of stimulated neurons might lead to a small change in overall engram connectivity. Although small, these decrements in connectivity accumulate over time, as a result of repetitive stimulation [41].

High turnover rates of synapses increase the volatility of network structure. This, in turn, poses a grand challenge to any synaptic theory of memory [42], and it is not yet clear how memories can at all persist in a system that is constantly rewiring [48]. In our model, the desired relative stability of memories is achieved by storing them with the help of a slow manifold mechanism. An estimation of turnover rates in our model amounts to about 18 % per day, which is comparable to the 20 % per day that have been measured in mouse barrel cortex [56]. In general, however, adult mice have more persistent synapses with much lower turnover rates as low as 4 % per month [40, 68, 28]. This can be accounted for in our model, as increased growth parameters of axonal, *β_a_*, and dendritic, *β_d_*, elements would lead to smaller synaptic turnover rates and, consequently, to more persistent spines (see Methods). The downside of increasing the growth parameters is that the learning process becomes slower. The turnover rate of 18% per day corresponds to a specific value of the parameter *β_d_*. It is conceivable, however, to implement an age-dependent parameter *β_d_*. For example, one could have a high turnover rate in the beginning and let the growth become slower with time. This would reflect the idea that the brains of younger animals are more plastic than the brains of older ones. As animals grow older, synapses become more persistent. Similar to certain machine learning strategies (“simulated annealing”), this could be the optimal strategy for an animal, which first explores a given environment and then exploits the acquired adaptations to thrive in it.

Recently, [14] showed that Hebbian structural plasticity could be the force behind memory consolidation through a process of stabilization of connectivity, which is based on the existence of an attractive fixed point in the plastic network structure. In our model, because of the decay along the slow manifold, memories are never permanent, and repeated stimulation is necessary to refresh them. We would argue, though, that forgetting is an important aspect of any biological system. In our case, we observe an exponential decay, if the stimulus is no longer presented. Furthermore, [4] has shown that a network can repair itself after lesion using a structural plasticity model similar to the one used in our current paper. Together with our results, this suggests that a structural perturbation of engrams (e.g. by removing connections or deleting neurons) could actually trigger a “healing” process and eventually rescue the memory. In the case of unspecific lesions, however, such perturbation might also lead to the formation of “fake” memories, or to the false association of actually unconnected memory items.

It appears that the attractor metaphor of persistent activity is not consistent with our model of homeostatic plasticity. As explained in Section 2.6, homeostatic control tends to delete connections between neurons which are persistently active, and in extreme cases could even lead to pathological oscillations. In the case discussed in Section 2.2, in contrast, the memories formed are “silent” (elsewhere classified as “transient” [52]), very different from the persistent activity usually considered in working memory tasks (elsewhere classified as “persistent”, or “dynamic” [52]). It was previously shown that silent memories can emerge in networks with both excitatory and inhibitory plasticity [61, 62, 47]. In all these cases, inhibitory plasticity allows the memory to be silent, but memory formation still relies on explicit Hebbian plasticity rules of excitatory connections. Some authors [62, 47] apply STDP to the excitatory connections, and others [61] just impose an increase in weight of excitatory-to-excitatory connections between neurons forming the assembly. As we understand it, such an increase could be achieved by Hebbian plasticity of excitatory connections. In contrast, our work shows that silent memories could potentially also be formed without correlation-based Hebbian plasticity on excitatory connections. Persistent activity, on the other hand, seems to be exclusively consistent with Hebbian plasticity models [14, 39, 64]. One consequence of the lack of persistent activity in our model is that pattern completion is restricted to the stimulation period. As previously seen, in our model, stimulating partial patterns leads to the subsequent activation of the full pattern (Figure 3C-D). Given that there is no persistent activity, however, the firing rates of all neurons go back to their baseline values as soon as the stimulus is gone. In models with persistent activity, on the other hand, a brief stimulation of partial patterns leads to activation of the full pattern, which remains active even after the stimulus is removed.

One possible way to integrate both mechanisms in a single network would be to keep their characteristic time scales separate. This could be accomplished, for example, by choosing faster time constants for Hebbian functional plasticity, and slower ones for homeostatic structural plasticity. An effective separation of time scales could also be obtained, if homeostatic structural plasticity would use somatic calcium as a signal, but not exert any control of the intermediate calcium levels in dendritic spines [25]. This might eliminate the need to specify a target rate in the model, and fast functional plasticity would shape connectivity in the allowed range of values where neurons have a distribution of firing rates reflecting previous experience. This induces a natural separation of time scales, where memories encoded by homeostatic plasticity would last much longer than in the present model, as only extreme transients would trigger rewiring. Homeostatic plasticity would perform Bayesian-like inference similar to structure learning, while functional plasticity would perform fast associative learning, similar to the system proposed by [19]. Integrating both functional and structural plasticity opens the possibility that different information is represented by the number of synapses and by the synaptic weights between pairs of neurons. This would allow for the encoding of more complex patterns. If the mean connectivity among neurons establishes the engram, the synaptic weights would offer additional degrees of freedom to encode information and to modulate the activity of individual neurons within the engram in an independent way. Different plasticity rules could thus be used to encode different temporal aspects of neuronal activity in either synaptic weights or synaptic connectivity.

Synaptic plasticity also influences the joint activity dynamics of neurons, which can be assessed with appropriate data analysis methods. Functional and effective connectivity, for example, are inferred from measured neuronal activity [20], [17], [34]. One implication of our results is that different activity dependent plasticity rules can lead to the same changes in functional and effective connectivity. Specifically, changes in network connectivity triggered by homeostatic plasticity also lead to changes in correlation. Therefore, these changes appear to be driven by correlation, and a correlation detector for each synapse might be postulated to implement it. This is, however, a wrong conclusion. The increased correlation and the Hebbian properties of the homeostatic model emerge as a network property due to availability of free elements and random self-organization. It remains a challenge to devise experiments that can differentiate between these fundamentally different possibilities.

We showed that very strong stimulation can damage the network by deleting too many synapses in a short time. The compensatory processes, which normally guarantee stability, get out of control and lead to seizure-like bursts of very high activity. This pathological behaviour of the overstimulated system could contribute to the etiology of certain brain diseases, such as epilepsy. The disruption of healthy stable activity is caused by a broken excitation-inhibition balance due to the high activity of one subgroup (Figure 7A). This, in turn, leads to the emergence of a runaway connectivity cycle (Figure 7E). Strategies for intervention in this case must take the whole cycle into account, and not just the phase of extreme activity. Inhibiting neurons during the high-activity phase, for example, could have an immediate effect, but it would not provide a sustainable solution to the problem of runaway connectivity. Our results suggest, against intuition maybe, that additional excitation of the highly active neurons could actually terminate the vicious cycle quite efficiently. It is important to note, however, that our system has not been designed as a model of epilepsy, and therefore does not reproduce all features of it [50]. In particular, seizure occurrence is stochastic in nature, but the limit cycle we describe here implies periodic activity and a periodic dynamics of connectivity. Although increased mossy-fibre connectivity among granule cells is known to be one of the structural changes related to epilepsy [3], there is currently no evidence for cyclic changes of this recurrent connectivity. Furthermore, the process of epileptogenesis in real brains is accompanied by other structural changes, such as neuronal death and glia-related tissue reorganization. In any case, our results shed light on a novel mechanism of pathological structural overcompensation and could potentially instruct alternative approaches in future epilepsy research.

Rene Descartes already proposed a theory of memory, paraphrased in [42]: Putting needles through a linen cloth would leave traces in the cloth that either stay open, or can more easily be opened again. Richard Semon, who originally coined the term “engram” in his book [51, 32], proposes that an engram is a “permanent record” formed after a stimulus impacts an “irritable substance”. Putting these two ideas together, we can think of Descartes’ needles not to penetrate an inanimate linen cloth, but a living brain, the irritable substance. In this case, we should expect the formation not of permanent holes, but of scar tissue, which grows further by repeating the procedure. Therefore, the memory of the system nothing else but a “scar” left by sensory experience. We think that this is a good metaphor for the type of memories described in our paper, formed through homeostatically controlled structural plasticity. Interestingly, this model was originally meant as a model for rewiring after lesion [4]. In our work, however, the “lesion” is imposed by stimulation, which induces phenomena similar to scar formation. The dynamics of this healing process is very universal, where resources from the whole network are used to fix a local problem leading to a scar. A perturbation introduces heterogeneity in a previously homogeneous organic substrate.

## 4 Methods

### 4.1 Network model

The neuronal network consists of *N_E_* = 10 000 excitatory and *N_I_* = 2 500 inhibitory current-based leaky integrate-and-fire (LIF) neurons. The sub-threshold dynamics of the membrane potential *V_i_* of neuron *i* obeys the differential equation

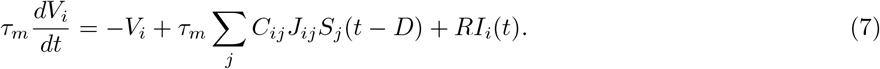

The membrane time constant *τ_m_* is the same for all neurons. The number of synaptic contacts between a presynaptic neuron *j* and a postsynaptic neuron i is denoted by *C_ij_*. The synaptic weights of individual contacts *J_ij_* is the peak amplitude of the postsynaptic potential and depends only on the type of the presynaptic neuron. Excitatory connections have a strength of *J_E_ = J* = 0.1 mV. Inhibitory connections are stronger by a factor *g* = 8 such that *J_I_* = –*g^J^* = —0.8mV. A spike train 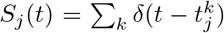 consists of all spikes produced by neuron *j*. The external input *Rl_i_*(*t*) to a given neuron in the network is conceived as a Poisson process of rate *v*_exl_ = 15 kHz and amplitude *τ_m_ J*. The external input to different neurons is assumed to be independent. All synapses have a constant transmission delay of *D* = 1.5 ms. When the membrane potential reaches the firing threshold *¼*_th_ = 20 mV, the neuron emits a spike that is transmitted to all postsynaptic neurons. Its membrane potential is then reset to *V_r_* = 10 mV and held there for a refractory period of *t*_ref_ = 2 ms.

The number of input synapses is fixed at 0.1*N_I_* for inhibitory-to-inhibitory and inhibitory-to-excitatory connections, and at 0.1NE for excitatory-to-inhibitory synapses. Once synaptic connections of these three types are established, they remain unchanged throughout the simulation. In contrast, excitatory-to-excitatory connections are initially absent and grow only under the control of a structural plasticity rule.

### 4.2 Plasticity model

Growth and decay of excitatory-to-excitatory (EE) connections follow a known model of structural plasticity regulated by firing rate homeostasis [4, 18]. In this model, each neuron *i* has a certain number of synaptic elements of two kinds available, axonal elements *α_i_*(*t*) and dendritic elements *d_i_*(*t*). These elements are bonded together to create functional synapses. Synaptic elements that have not yet found a counterpart are called free elements, denoted by 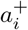 and 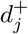, respectively. If 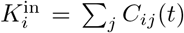 denotes the in-degree and 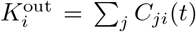 the out-degree of neuron *i*, the number of free elements in every moment is given by 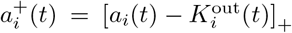 and 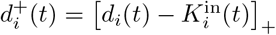, with [*x*]_+_ = max(*x*, 0).

Firing rate homeostasis is implemented by allowing each neuron to individually control the number of its synaptic elements. We assume that each neuron *i* maintains a time-dependent estimate of its own firing rate, using its intracellular calcium concentration *φ_i_*(*t*) as a proxy. This variable reflects the spikes *S_i_* (*t*) the neuron has generated in the past, according to

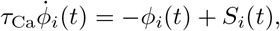

with a time constant of *τ*_Ca_ = 10 s for all neurons. This implements a first-order low-pass filter. In this model, more weight is given to more recent spikes, following a decaying exponential. The calcium trace *φ_i_*(*t*) of individual neurons is used as a control signal for the number of axonal elements *a_i_*(*t*) and dendritic elements *d_i_*(*t*) according to the homeostatic equations

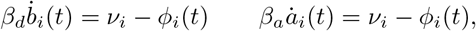

where *β_d_* is the dendritic and *β_a_* is the axonal growth parameter. Both have a value of 2 in our default setup. The parameter *ν_i_* is called the target rate. Whenever the firing rate estimate (in fact, the calcium concentration) is below the target rate, the neuron creates new axonal and dendritic elements, from which new synapses can be formed. Whenever the estimated firing rate is larger than the target rate, the neuron deletes some of its elements, removing the synapses they form. The (negative) decrements 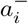 and 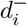 in the number of synaptic elements, respectively, are in each moment given by 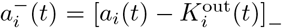 and 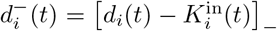 for [*x*]_ = min(*x*, 0).

After time intervals of duration Δ*T_s_*, all negative elements are collected, and 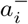 of the existing outgoing connections of neuron *i* are randomly deleted. Similarly 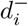 connections are randomly deleted. The deletion of bonded elements of one type frees their counterparts that were previously connected to the deleted element. Then, all free dendritic |**d**^+^| and axonal elements |**a**^+^| are collected and randomly combined into pairs, creating *n* = min(|**a**^+^|, |**d**^+^|) new synaptic connections. This algorithm has originally been devised by [4], an efficient implementation of it in NEST exists [12] and has been employed for all our simulations.

### 4.3 Mathematical re-formulation of the algorithm

The algorithm of homeostatically controlled structural plasticity can be expressed as a discrete-time stochastic process. Rewiring takes place at regular intervals of duration Δ*T_s_*

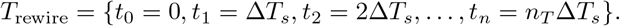

Between any two rewiring events, for *t* ∈ (*t_k_, t_k_* + Δ*T_s_*), the neuron just accumulates synaptic elements

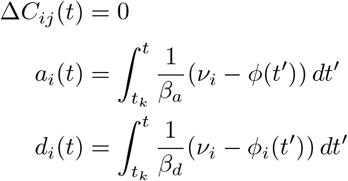

while the already established connectivity remains unchanged. At every rewiring step, the rearrangement of connectivity is completely random, driven by the probabilities *P*(Δ_±_*C_ij_*(*t_k_*) = *c*|**a**(*t_k_*), **d**(*t_k_*), **C**(*t*_*k*_1_)) of creating or deleting c connections from neuron *j* to neuron *i* at time *t_k_*, during a time step of duration Δ*T_s_*. This gives rise to a discrete-time stochastic process

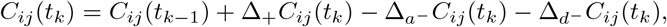

for *t_k_* ∈ *T*_rewire_. Here, we define Δ_*a*_. *C_ij_*(*t_k_*) as the random variable representing the creation of synapses, while Δ_*a*_-*C_ij_* (*t*) and Δ_*d*_-*C_ij_* (*t*) are random variables describing the deletion of synapses by removing their corresponding axonal and dendritic elements, respectively.

We first calculate the probability to create just one new connection 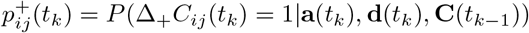. We can express this as a process of selecting a presynaptic partner *j* with probability 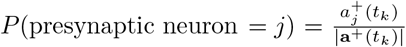 and then a postsynaptic partner *i* with probability 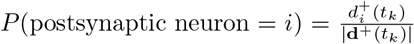. We then connect the pair with the product of both probabilities

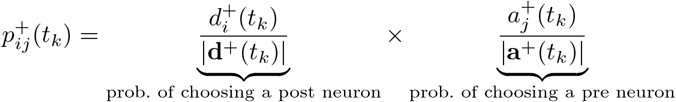

as they are independent random variables. We now have the probability of creating individual connections, but the full probability of an increment for the whole network is hard to calculate. This is because of statistical dependencies that arise from the fact that rewiring affects all neurons in the network simultaneously. First, the total number of new connections is *n* = min(|**a**^+^|, |**d**^+^|). Second, the number of new connections for a given pair of neurons is bounded by the number of free axonal and dendritic elements in the two neurons, respectively, 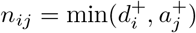. Finally, we cannot delete more connections than we actually have. This indicates that independent combinations of individual probabilities cannot be expressed by simple binomial distributions, but a hypergeometric distribution arises instead. We obtain for the probability of creating *c* synapses from neuron *j* to neuron *i*

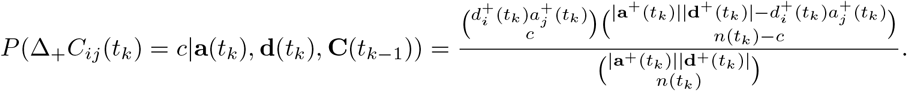

This probability is easy to understand: We divide the ensemble of all possible new synapses (which has size |**a**^+^ ||**d**^+^1) into the ensemble of potential synapses between the pair (*i, j*) (which has size 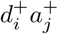) and all the rest. We then choose *c_ij_* < *n_ij_* (*t_k_*) connections from the preferred ensemble, and the rest of connections from the remaining pool. This is “sampling without replacement” as there is a fixed number of new connections *n*(*t_k_*) in each time step.

We can now calculate the probability to delete one connection using axonal “negative” elements as 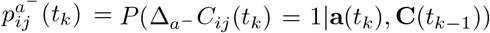. This is the probability to choose one to-be-deleted element from all existing elements 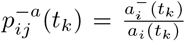 and 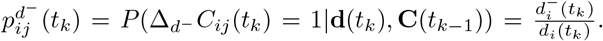 Out of *α_i_*(*t_k_*) candidates for deletion, we select 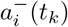, subject to the condition not to delete more than *C_ij_* (*t_k_*) for this particular pair of neurons. This constraint is reflected by the hypergeometric distribution. The preferred population is represented by the elements bonded into connections from neurons *j* to neuron *i*, and the other population is comprised by all remaining elements of neuron *j*. Finally, during rewiring events in *T*_rewlre_, we obtain for the stochastic evolution of *C_ij_* (*t_k_*)

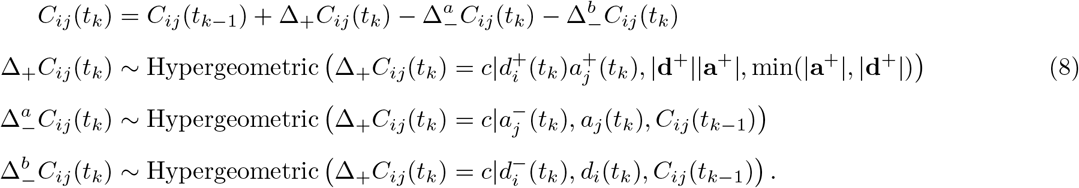

To calculate the total change Δ*C_ij_*(*t_k_*) we have to account for all these contributions. The distribution of the total increment is not simply the convolution of the three distributions given above, as they are not independent. On the contrary, (negative) decrements occasionally influence (positive) increments, as we first delete connections and thereby create additional free elements. Therefore, the number of free elements has to be corrected as

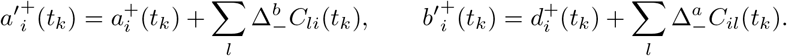

Equation 8 defines a complicated discrete-time stochastic process, but since we are at this point interested only in the expected change of connectivity, we can restrict ourselves to the evolution of expectations. We use 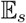 to denote the (linear) operator of expectation over the structural noise, i.e. over the realizations of the increments/decrements Δ*C_ij_* (*t_k_*). We have

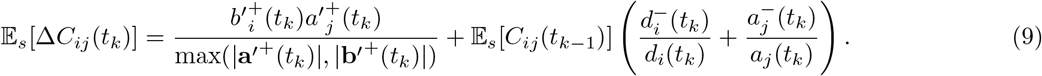

Further on in this paper, for notational convenience, we write 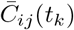 for the expectation 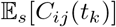.

### 4.4 Time-continuous limit

We now switch over to continuous equations, which result from the limit Δ*T_s_* → 0. In this case, we can express free elements and negative elements as

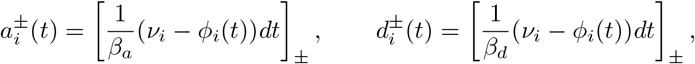

which will assign infinitesimally small values to the numbers of free elements and negative elements. The bracket notation [*x*]_±_ corresponds to flooring/ceiling operations using Heaviside step function [*x*]_±_ = *θ*[± *x*]. Since the rewiring takes place continuously, the numbers of elements are, up to infinitesimal correction, the same as the degrees, 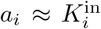 and 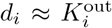. We can now define the rates of creation and deletion of axonal elements as 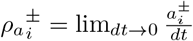 and dendritic of elements as 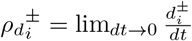 and we write explicitly

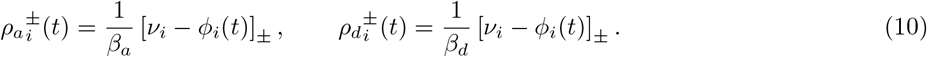

As noted above, we have omitted here the expectation over the structural noise 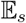 from the notation. We can now calculate the evolution of connectivity from Equation 9 as

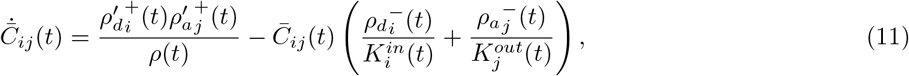

where 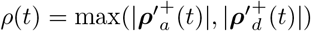. The specific implementation requires that the deletion of elements takes place first, with the corresponding synaptic partner remaining available as a free element to form new connections. After an axonal element has been deleted, the previously bonded dendritic element becomes a free element. This requires a correction on the rate of free elements

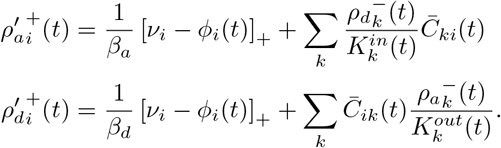

### 4.5 Spiking noise

We assume that spike trains 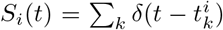 reflect an asynchronous-irregular state of the network, so they can be modeled as a stochastic point process. As described in [2, 58], the coefficient of variation of the spike trains generated by a leaky integrate-and-fire neuron driven by Gaussian white noise current is given by

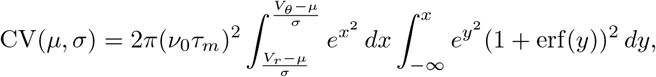

where *μ* and *σ* are the parameters of the current input, and *τ_m_*, *V_r_* and *θ* are the parameters of the neuron, respectively. The configuration used here yields CV ≈ 0.7. A good approximation for spike trains of a given rate *r* and irregularity CV is obtained with a specific class of renewal processes, so-called Gamma processes. These have an ISI distribution 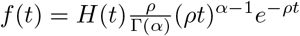, with parameters 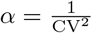 and 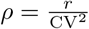.

In our model, the fluctuating intracellular calcium concentration is a shotnoise, a continuous signal that arises from a point process through filtering. Here, the point process has a mean rate *ν*, and the calcium signal is a convolution with an exponential kernel 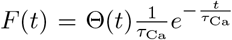. Campbell’s theorem allows us compute the mean 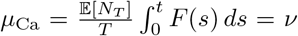 and the variance 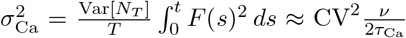 of the calcium variable. Here we used the fact that spike count of Gamma process 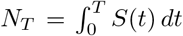 has a mean 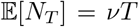 and a variance 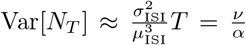, provided the observation time T is long enough [46]. As a consequence of the Central Limit Theorem, the amplitude distribution of the calcium signal is approximately Gaussian 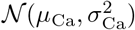, provided the mean spike rate is much larger than the inverse time constant of the calcium signal 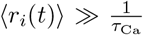. In other words, if the mean firing rate is 8 Hz and the calcium constant is *τ*_Ca_ = 10 s, there are on average 80 spikes in the characteristic time interval *τ*_Ca_.

We are actually interested in the equilibrium rates of free and negative elements in Equation 10. These rates are rectified versions of the shifted calcium trace, and in order to calculate them we resort to the ergodic theorem and to an adiabatic approximation. The former is a self-averaging property, according to which the time-averaged variable for a long observation is equal to the equilibrium mean value 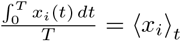. The adiabatic property comes from the fact that spiking dynamics of a network is much faster than network remodeling due to structural plasticity. At every point in time, plasticity is driven by the average adiabatic rate. This implies that, at every point in time *t*, the structural plasticity sees the equilibrium distribution of the calcium trace 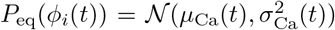. The right-hand side of Equation 10 becomes

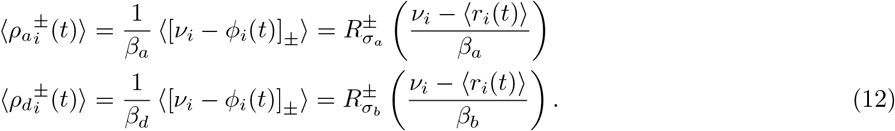

The transfer function for synaptic elements is given by 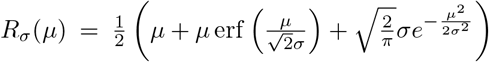, and the variance of the rate of elements is 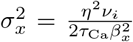. The parameter *η* is a correction factor, which accounts for the regularity of spike trains. In our case *η* = CV. Even if the mean number of free elements is zero, the noise will still drive the creation and deletion of elements with rate 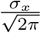. Even if homeostatic control manages to drive all neurons to their target rates, the inherent calcium fluctuations and the associated uncertainty of firing rate inference will still induce random rewiring.

### 4.6 Mean-field approximation of population dynamics

We define population means for variables *x* ∈ {*r, φ, α, b*} and neuronal populations *Y, Z* as

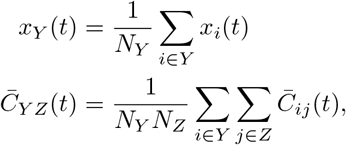

where *N_Y_* is the size of population *Y*. An individual neuron in population *Y* typically gets many inputs from every other neuronal population, which invites use of mean-field approximation due to the central limit theorem. The resulting currents aggregate to a Gaussian white noise process with mean and variances given as

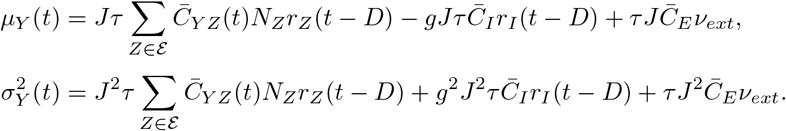

The stationary firing rate of a leaky integrate-and-fire neuron driven by input with mean *μ* and variance *σ*^2^ is [2]

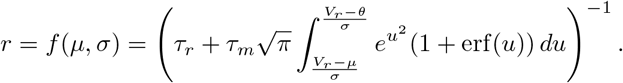

Solving the time-dependent self-consistency problem for multiple interacting plastic populations is a challenging problem. We suggest here to use an adiabatic approximation, resorting to the fact that the firing rate dynamics is much faster than the plastic growth processes. Therefore, we employ a Wilson-Cowan type of the firing rate dynamics [36]

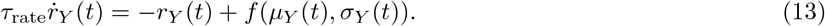

The relaxation time is set to *τ*_rate_ < *τ_m_* to account for the fact that population response is generally much faster than the membrane potential dynamics [21]. The parameter *τ*_rate_ is the only free parameter in this model, but our results do not depend on its exact value as long as *τ*_rate_ ≤ *τ_m_*. The heuristic described by Equation 13 results in a tractable and numerically stable system to be analyzed with standard dynamical system tools.

As the equation is linear, the average calcium trace can be computed without approximation

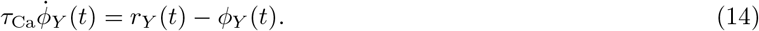

The rate of element creation and deletion is also linear, therefore population averages are given by

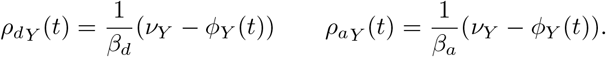

Finally we calculate the average connectivity

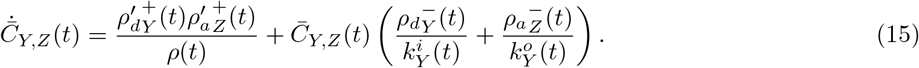

Here,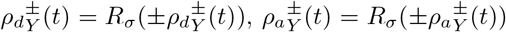, and *ρ*(*t*) is same as before, while the corrected rate for free elements is

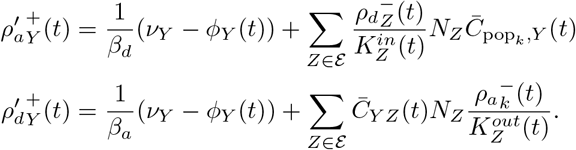

### 4.7 Line attractor of the deterministic system

We represent the state of the network consisting of one static inhibitory and two plastic excitatory ensembles as a vector

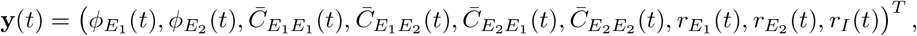

the components of which adhere to the calcium dynamics Equation 14, the connectivity dynamics Equation 15, and the activity dynamics Equation 13. The joint ODE system 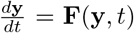 defines the vector field **F**. We first explore the stationary states of the deterministic system (*σ_x_* = 0), setting the left hand side of all ODEs to zero. In this case, calcium concentration and excitatory firing rates are fixed at their target values, and inhibitory firing rates 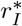 can be obtained using the self-consistency Equation 13. Stationary connectivity is calculated from the condition that the rates of creation and deletion of synaptic elements are zero. This implies that the numbers of free axonal and dendritic elements are zero, **a**^±^ = **0** and **d**^±^ = **0**, which in turn implies that **a** = **1**^*T*^ · **C** and **d** = **C** · **1**. The second condition is that all axonal elements are bonded with a dendritic element, and the total number of both types of elements are the same |**a**| = |**d**|.

Let us consider the case of two plastic excitatory ensembles *E*_1_ and *E*_2_ and one static inhibitory population *I*. Using the parameter 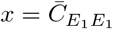, the stationary state of the deterministic network is a line

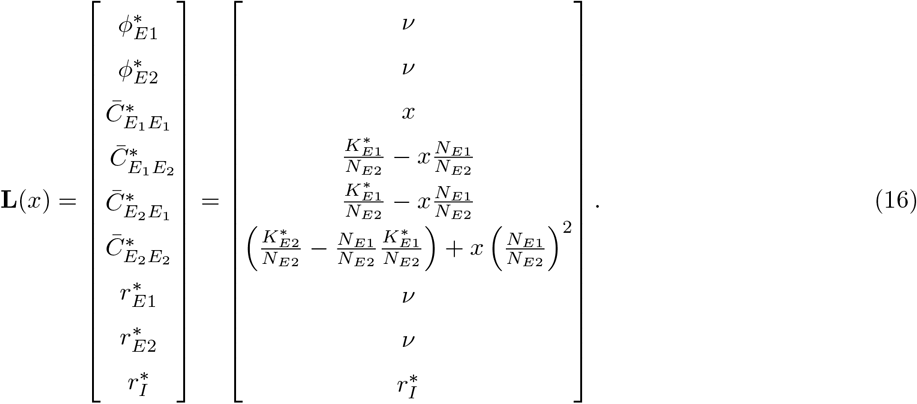

Here, 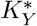 is the mean stationary in-degree, and we assume that the dendritic element growth factor *β_d_* is less or equal to the axonal element growth factor *β_a_*. The attractor is a line segment, as a consequence of linear conditions for element numbers and non-negativity of excitatory connections 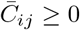. This solution can be easily generalized to a solution for individual connections of NE excitation neurons, or for the case of *N_E_* excitatory populations. The minimal invariant changes of the stationary connectivity matrix **C** that keep in-degree and out-degree conditions valid are of the type

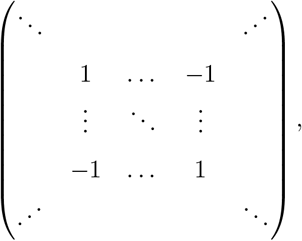

where all missing entries are considered to be zero. This transformation defines a hyperplane section invariant space. The stationary connectivity can be solved as

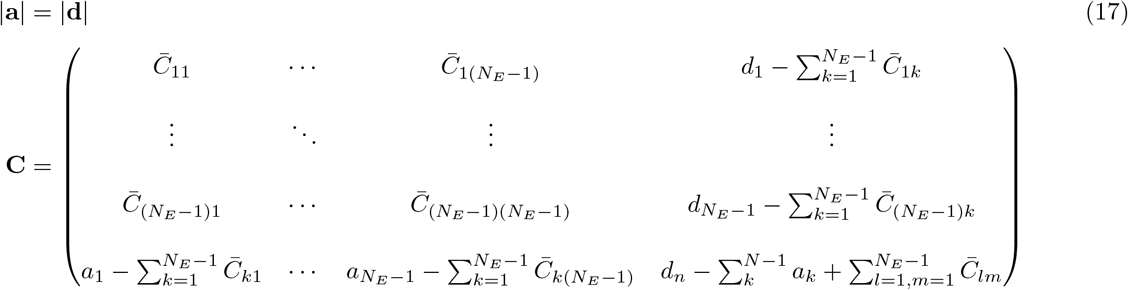

Note that we have put the in-degree conditions in the last row and the out-degree conditions in the last column, but there is a permutation symmetry and they could apply to any row-column combination. The hyperplane for individual neurons has *N_E_*(*N_E_* — 3) + 1 dimensions, if there are no self-connections. In the case of *n_E_* excitatory populations, the hyperplane has *n_E_* (*n_E_* — 2) + 1 dimensions.

### 4.8 Slow manifold and diffusion to the global fixed point

Now we calculate the stationary state for the stochastic system and assess the geometry and stability of the underlying phase space. In the case of finite noise, *σ_d_* > 0, Equation 11 has an attractive fixed point, instead of a hyperplane attractor. The fixed point is given by

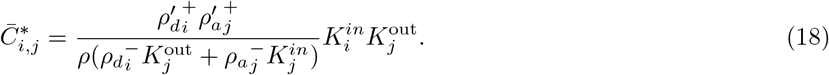

Here, without loss of generality, we have assumed that the growth rate of dendritic elements is smaller or equal as compared to axonal elements. We obtain a normalized outer product of the indegree and outdegree vectors, respectively. In the case of identical dynamics for axonal and dendritic elements, and if all neurons have the same target rate *v_i_* = *v*, we obtain the fixed point by plugging in the homogeneous solution

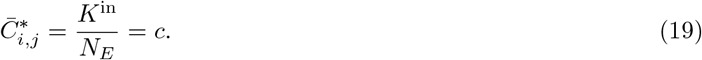

This fixed point depends only on the indegree of neurons. For a two-population system, the fixed point is 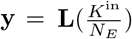, the most entropic configuration of the line segment attractor. This suggests a special importance for the relic of the line-attractor, which, as we show later, turns into a stochastic slow manifold through the presence of noise.

We now calculate the relaxation time for the three-population system to assess the persistence of a memory trace. To this end, we calculate the Jacobian 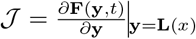 on the slow manifold, using a linearization of the vector field **F** (see Section 4.7). For the stochastic system, the Jacobian 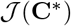 about the global fixed point is

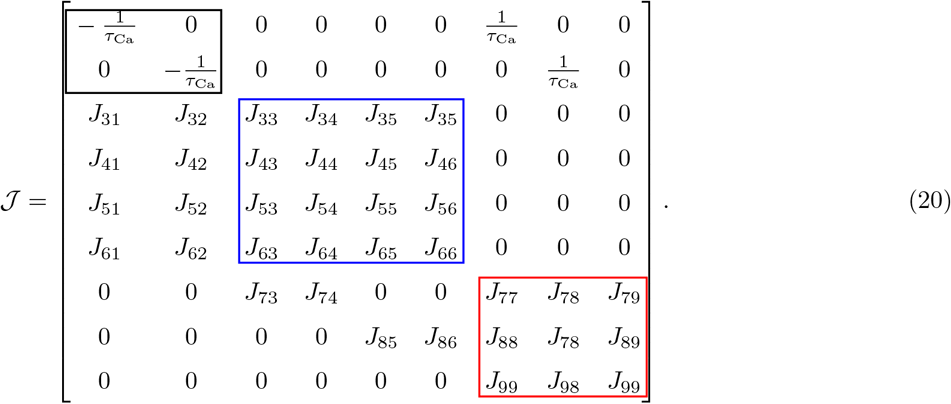

The Jacobian has a block structure, which corresponds to calcium activity (black rectangle in Equation 20), connectivity 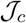 (blue rectangle) and spike activity 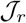 (red rectangle). The connectivity block around fixed point **y** = **L**(*c*) is given by

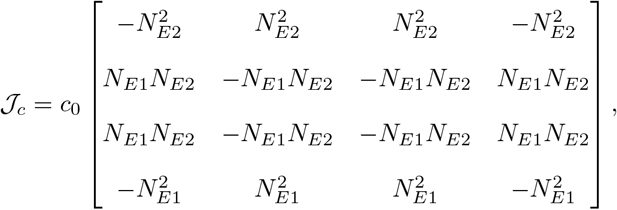

where 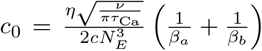. Jacobian entries responsible for interaction between connectivity variables and calcium activity variables are 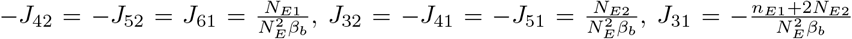 and 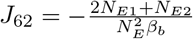. The spike activity terms in the Jacobian are

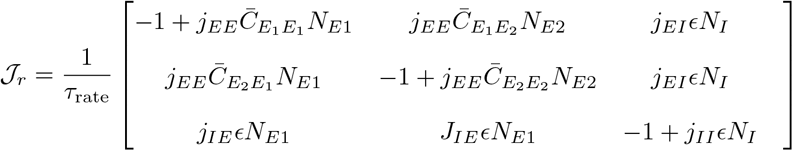

Here we define the effective interaction of different neuron types neurons as excitatory-to-excitatory 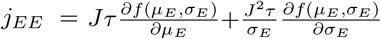, inhibitory-to-excitatory 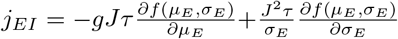, excitatory-to-inhibitory 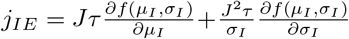 and inhibitory-to-inhibitory neurons as 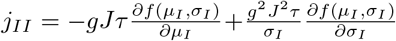. The Jacobian elements which are responsible for influence of connectivity change to rates are 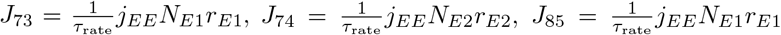 and 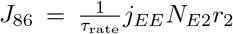. These terms are the largest in the Jacobian, but since connectivity changes only through a change of calcium, they will not play an essential role for long-term stability.

We exploit the block structure of the Jacobian 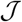 to separate between connectivity variables, the fast firing rate and calcium variables. We can solve the connectivity eigenproblem analytically,

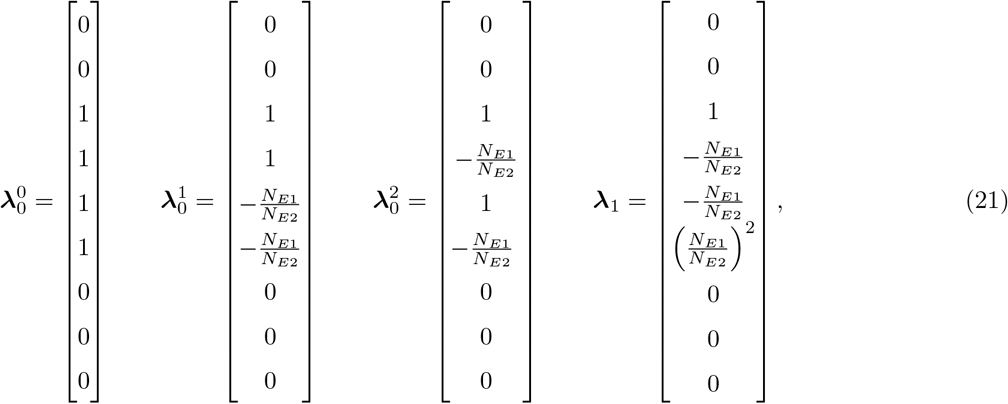

with eigenvalues λ_0_ =0 and 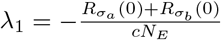.

The persistence of a memory trace in the system with noise is essentially determined by the diffusion to the global fixed point. In fact, the line attractor of the noiseless system turns into a slow manifold of the noisy system. The eigenvalue λ_0_ is degenerate, and the three-dimensional invariant space defines the central manifold of the connectivity subsystem. The eigenvalue λ_1_ is also an eigenvalue of the full Jacobian 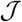 and it represents the slowest time scale of the system in the direction of the slow manifold. It is easy to check that 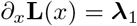. This means that the noiseless line attractor is exactly corresponding to the slow manifold of the system. The other eigenvectors have components orthogonal to the slow manifold. Those are dominated by fast variables, and they relax quickly. As a result, the relaxation dynamics is dominated by the eigenvalue λ_1_, and the relaxation time of the system under consideration is

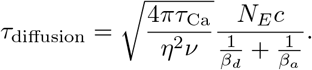

For the default parameters used here, its value is around 5 000 s, fully in accordance with numerical simulations. The system relaxes along the direction of the slow manifold. Therefore, although rates of creation and deletion are identical on the slow manifold, the system takes a fast trajectory off the slow manifold upon perturbation, and it relaxes slowly along the slow manifold towards the fixed point.

A quantitative measure for the volatility of network structure used in experiments is the turnover ratio (TOR) of dendritic spines [56]. It is defined as 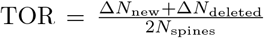, where the changes Δ*N*_new_ and Δ*N*_deleted_ are typically measured per day. For the case when the network plasticity is selectively driven by diffusion, this quantity corresponds to the eigenvalue λ_1_, corresponding to a value around 18% per day for standard parameters.

### 4.9 Linear stability analysis

Numerical exploration of the eigenvalues of the Jacobian 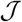 (Equation 20) show that for our system there is linear stability for a wide parameter range. However this system can produce damped oscillations, which may cause problems. In the case discussed here, dendritic and axonal elements have identical dynamics, *β_a_* = *β_d_*, and damped oscillations influence the connectivity block in the same way. Therefore it was sufficient to consider a reduced system of only one plastic excitatory population and one static inhibitory population to analyze the influence of the calcium time constant and element growth. This gives rise to the Jacobian

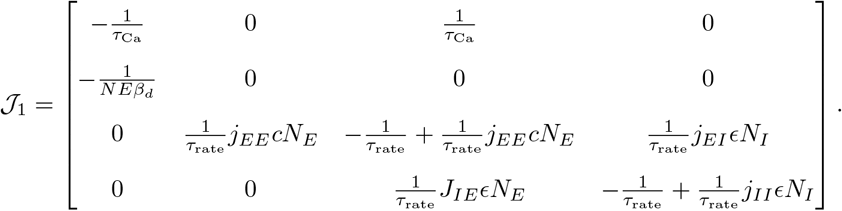

The eigenvalues of 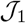 have been reduced to radicals using *Mathematica* 12.0, and the complex conjug values λ_3/4_ responsible for oscillations are plotted in Figure 6.

Although there is linear stability around the global fixed points for wide parameter range, the same true for all points along the line attractor. The connectivity eigenvalue λ_1_ is constant on the slow mani one population presents large recurrent connectivity, spike activity jumps to a persistent high activity can track this instability in the Jacobian 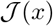 along the line attractor **L**(*x*). As the recurrent connect] first ensemble *E*_1_ is increased, one of the eigenvalues becomes positive, and its corresponding eigenve the biggest contributions in the direction of spike activity *r*_*E*_1__ and *r*_*E*_2__. Here we use this fact and use *t* Jacobian

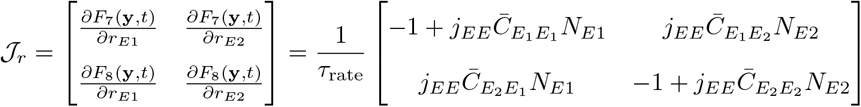

to find the approximate position of this transition on the line attractor. The spiking activity becomes unstable, when the real part of the second eigenvalue 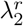 becomes positive, and at this point the determinant of the Jacobian changes its sign (since both eigenvalues are real). We can use this criterion to determine when the system loses linear stability, leading to the critical value *c*_crit_ of connectivity 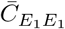

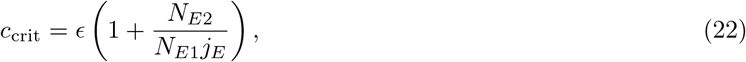

where *j_E_* is a dimension-less measure of effective excitatory excitability *j_E_* = *ϵN_EjEE_*. The value *c*_crit_ is the upper bound of full stability, but the system loses stability even below this value, as discussed in our results.

### 4.10 Network simulations

All simulations have been performed using the neural network simulator NEST 2.16.0 [37].

#### 4.10.1 Grown networks

All numerical stimulation experiments start from grown networks. To this end, we initialize a network with random connections to and from inhibitory neurons and no excitatory-to-excitatory connections, whatsoever. The latter are then grown under the control of homeostatic structural plasticity. The controlled variable is the firing rate of excitatory neurons, the target rate is set to *ν* = 8 Hz for all neurons. During this initial growth period, excitatory neurons receive external Poisson input of rate *v*_ext_ = 15 kHz. After a long-enough growth time *t*_growth_, the network structure has reached its equilibrium and all neurons fire at their target rate, apart from small fluctuations.

#### 4.10.2 Conditioning paradigm

Non-overlapping neuronal ensembles comprising 10 % of all excitatory neurons each are selected and labeled US, C1 and C2. Non-plastic connections of weight *J_E_* = 0.1 mV are created from all neurons labeled US to a readout neuron, which has the same properties as all the other neurons in the network. Stimulation is specific for a certain group of neurons. A stimulation cycle consists of an increased external input rate of 1.4 *v*_ext_ for a period of 2s, followed by a relaxation period of 48 s, during which the external input is set back to *v*_ext_.

The whole protocol consists of 5 different episodes: growth, baseline, encoding, decay and retrieval. During “baseline” each of the 3 groups is stimulated alone in the order US, C1, C2. The “encoding” episode consists of 6 stimulation cycles. In 3 out of the 6 cycles, C2 is stimulated alone. In the other 3 cycles, neurons from both US and C1 are stimulated at the same time. The “decay” episode lasts 100 s, during which no stimulation beyond *v*_ext_ is applied. During “retrieval”, there are 2 stimulation cycles, C1 alone is followed by C2 alone. Connectivity is recorded every 1 s. In this protocol, plasticity is always on, and all measurements are performed in the plastic network. Growth time is *t*_growth_ = 100 s, the remaining parameters are *β_d_* = *β_a_* = 0.4, *τ*_Ca_ = 1 s, Δ*T_s_* = 10 ms.

#### 4.10.3 Repeated stimulation

Starting from a grown network, a random neuronal ensemble comprising 10 % of all excitatory neurons in the network is repeatedly stimulated for 8 cycles. Here, a stimulation cycle consists of a stimulation period of 150 s, during which the external input to the ensemble neurons is increased to 1.05 *v*_ext_. It is followed by a relaxation period of 150 s, during which the external input is set back to *v*_ext_. During stimulation, the connectivity is recorded every 15 s. After encoding the engram, plasticity is turned off, and all measurements are now performed in a non-plastic network. Growth time is *t*_growth_ = 500 s, the remaining parameters are *β_d_* = *β_a_* = 2, *τ*_Ca_ = 10 s, Δ*T_s_* = 100 ms.

#### 4.10.4 Readout neuron

Two non-overlapping ensembles comprising 10 % of all excitatory neurons each are randomly selected and labeled ¦ *A*_1_ and *A*_2_. Starting from a grown network, *A*_1_ is stimulated twice. During each stimulation cycle, the external input to stimulated neurons is increased to 1.1 *v*_ext_ for a time period of 150 s. This is followed by a relaxation period of 150 s, during which the external input is set back to *v*_ext_. After a pause of 100 s, *A*_2_ is stimulated once using otherwise the same protocol. Growth time is *t*_growth_ = 500 s, the remaining parameters are *β_d_* = *β_α_* = 2, *τ*_Ca_ = 10 s, Δ*T_s_* = 100 ms.

After the encoding of engrams, plasticity is turned off. A readout neuron is added to the network, which has the same properties as all other neurons in the network. Non-plastic connections of weight *J_E_* = 0.1 mV are created, from a random sample comprising 9 % of all excitatory and 9 % of all inhibitory neurons in the network. Two i new non-overlapping ensembles comprising 10 % of all excitatory neurons each are selected as random patterns *A*_3_ and *A*_4_. Neurons in the network are stimulated in the order *A*_3_, *A*_2_, *A*_1_, *A*_4_. During each stimulation cycle, the external input to neurons in the corresponding group is increased to 1.1 *v*_ext_ for a period of 1 s duration. This is i followed by a non-stimulation period of 4 s, during which external input rate is set back to *v*_ext_.

#### 4.10.5 Formation and decay of engrams

Starting from a grown network, a subgroup comprising 10 % of all excitatory neurons is randomly selected and stimulated for 150 s, followed by a prolonged relaxation period of 5 500 s. During stimulation, the external input to stimulated neurons is increased to 1.1 *v*_ext_. For all simulations, *β_d_ = β_a_* = 2, Δ*T_s_* = 100 ms.

The simulations to demonstrate how *τ*_decay_ changes with *τ*_Ca_ were performed with a target rate *ν* = 8 Hz and *τ*_Ca_ = 2,4, 8,16, 32 s. The simulations showing how *τ*_decay_ changes with *v* were performed with *τ*_Ca_ = 10 s and *v* = 2,4, 8,16, 32 Hz. In the simulations performed with *ν* = 2, 4 Hz, *τ*_growth_ = 5000 s, neurons are stimulated for 1 500 s, and the relaxation period is 26 000 s. The parameter *τ*_decay_ is estimated from simulated time series by performing a least-squares fit of an exponential function to the connectivity values during the decay period, and extracting the fitted time constant.

#### 4.10.6 Linear stability

Starting from a grown network, a subgroup comprising 10 % of all excitatory neurons is randomly selected and stimulated for a time period *t*_stlm_. During stimulation, the external input to stimulated neurons is increased to 1.1 *v*_ext_. Stimulation is followed by a period of duration *t*_relax_, during which the external input is set back to *v*_ext_ and the connectivity relaxes back to a new equilibrium. The parameters used are for non-oscillatory, regime: *t*_growth_ = *t*_stim_ = *t*_relax_ = 800 s, *β_d_* = *β_a_* = 2, *τ*_Ca_ = 5s, Δ*T_s_* = 1ms; for the weakly oscillatory regime: *t*_growth_ = *t*_stim_ = *t*_relax_ = 400 s, *β_d_* = *β_a_* = 0.2, *τ*_Ca_ = 20 s, Δ*T_s_* = 0.2 ms; and for the strongly oscillatory regime: *t*_growth_ = *t*_stim_ = *t*_relax_ = 200 s, *β_d_* = *β*_a_ = 0.03, *τ*_Ca_ = 10 s, Δ*T_s_* = 0.2ms.

#### 4.10.7 Non-Linear stability

Starting from a grown network, a subgroup comprising 10% of all excitatory neurons is randomly selected and stimulated for 200 s. During stimulation, the external input rate to stimulated neurons is increased to 1.25 *v*_ext_. Stimulation is followed by a relaxation time in which external input rate is set back to *v*_ext_. Growth time *t*_growth_ = 500 s, *β_d_* = *β_α_* = 2, *τ*_Ca_ = 10 s, Δ*T_s_* = 0.1 ms. During and after stimulation, connectivity is recorded every 5s.

#### 4.10.8 Overlap measure

The similarity between network activity during stimulation, spontaneous and evoked responses is measured by the corresponding overlaps [22] defined as

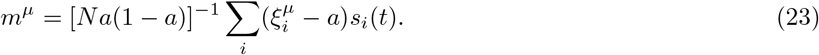

The pattern *ξ^μ^* is a vector of dimension *N*. Each entry 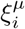 is a binary variable indicating whether or not neuron *i* is stimulated by pattern *ξ^μ^*, and it has a mean value of 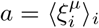. The activity vector *s_i_*(*t*) is also composed of binary variables, which indicate whether or not neuron *i* is active in a given time bin. In all figures, the bin size used for calculating overlaps is 10 ms.

#### 4.10.9 Population firing rate

In Figure 3B, the population response of excitatory neurons is estimated for different connectivity values using a mean-field rate model [2]. We consider a model with three populations, two of which are excitatory (*E*_1_ comprises 10% and *E*_2_ comprises 90% of all *N_E_* = 10 000 excitatory neurons), and one is inhibitory with *N_I_* = 2 500 neurons. All connectivities involving inhibitory neurons are fixed and set to *ϵ* = 0.1. For the excitatory-to-excitatory connections, we systematically vary the connectivity within the *E*_1_ population 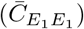, and calculate the other values to achieve a constant excitatory in-degree of *ϵN_E_*. All other parameters used are unchanged. For the different values of 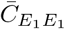 we calculate the population rate of excitatory neurons for *E*_1_ receiving a larger external input of 1.05 *v*_ext_ as compared to the *E*_2_ and *I* populations (*v*_ext_ = 15 kHz).

#### 4.10.10 Pattern completion

To address pattern completion, we employ the non-plastic network after engram encoding (see 4.10.3). Different fractions of the neurons belonging to the engram are stimulated for 10 s, and we calculate the overlap averaged over the stimulation time 〈*m*^*E*_1_^). For each fraction of stimulated neurons, 50 different simulations are run, during which a different subsample of the engram neurons is stimulated.

**Table 1:** List of symbols.

| Symbol | Description |
| --- | --- |
| $\phi(t)$ | Calcium trace |
| $\tau_{\text{Ca}}$ | Calcium time constant |
| $S(t)$ | Spike train |
| $r(t)$ | Instantaneous firing rate |
| $\nu$ | Target rate |
| $a(t)$ | Number of axonal elements |
| $d(t)$ | Number of dendritic elements |
| $\beta_d$ | Dendritic growth parameter |
| $\beta_a$ | Axonal growth parameter |
| $C_{ij}(t)$ | Number of synaptic connections from neuron $j$ (presynaptic) to neuron $i$ (postsynaptic) |
| $\bar{C}_{ij}(t)$ | Expected value of number of synaptic connections $C_{ij}(t)$ |
| $E$ | Excitatory neurons |
| $E_1$ | Stimulated excitatory neurons |
| $E_2$ | Non-stimulated excitatory neurons |
| $I$ | Inhibitory neurons |
| $\tau_{\text{drift}}$ | Effective time constant of encoding |
| $\tau_{\text{diffusion}}$ | Effective time constant of forgetting |
| $m^x(t)$ | Overlap of network activity with pattern $x$ |
| $\tau_{\text{rate}}$ | Relaxation time of rate dynamics |

## 5 Acknowledgements

Supported by Erasmus Mundus / EuroSPIN, DFG (grant EXC 1086) and Carl Zeiss Foundation. The HPC facilities are funded by the state of Baden-Württemberg through bwHPC and DFG grant INST 39/963-1 FUGG. We thank Sandra Diaz-Pier and Mikaël Naveau from the Research Center Jülich for support on new features of NEST, and Uwe Grauer from the Bernstein Center Freiburg as well as Bernd Wiebelt and Michael Janczyk from the Freiburg University Computing Center for their assistance with HPC applications.

## Notes

### Competing Interest Statement

The authors have declared no competing interest.

